# DNAJC13 localization to endosomes is opposed by its J domain and its disordered C- terminal tail

**DOI:** 10.1101/2024.12.19.629517

**Authors:** Hayden Adoff, Brandon Novy, Emily Holland, Braden T Lobingier

## Abstract

Endosomes are a central sorting hub for membrane cargos. DNAJC13/RME-8 plays a critical role in endosomal trafficking by regulating the endosomal recycling or degradative pathways. DNAJC13 localizes to endosomes through its N-terminal Plekstrin Homology (PH)-like domain, which directly binds endosomal phosphoinositol-3-phosphate (PI(3)P). However, little is known about how DNAJC13 localization is regulated. Here, we show that two regions within DNAJC13, its J domain and disordered C-terminal tail, act as negative regulators of its PH-like domain. Using a structure-function approach combined with quantitative proteomics, we mapped these control points to a conserved YLT motif in the C-terminal tail as well as the catalytic HPD triad in its J domain. Mutation of either motif enhanced DNAJC13 endosomal localization in cells and increased binding to PI(3)P *in vitro*. Further, these effects required the N-terminal PH-like domain. We show that, similar to other PI(3)P binding domains, the N-terminal PH-like domain binds PI(3)P weakly in isolation and requires oligomerization for efficient PI(3)P binding and endosomal localization. Together, these results demonstrate that interaction between DNAJC13 and PI(3)P serves as a molecular control point for regulating DNAJC13 localization to endosomes.

**Significance Statement:**

- DNAJC13 controls endosomal sorting by regulating proteins which mediate the endosomal recycling and degradative subdomains.
- Here we show that subcellular localization of DNAJC13 is regulated through the coordinated action of three of its domains: the PH-like domain which has low affinity for PI(3)P, the J domain, and a YLT motif in its disordered C-terminus.
- This study defines a novel mechanism by which DNAJC13 function is regulated.

## Introduction

Endosomes function as critical sorting hubs in the cell where membrane proteins are selectively sorted for degradation at the lysosome or for recycling to the Golgi or plasma membrane (Cullen and Steinberg, 2018). To achieve this function, endosomes host multiple proteins and protein complexes—spatially restricted into degradative and recycling domains— which select membrane protein cargos for distinct destinations (Cullen and Steinberg, 2018). The recycling subdomain is marked by proteins which assist in removal of proteins from the maturing endosomal system, including sorting nexins like SNX1, the Retromer complex, and the actin nucleating WASH complex (Cullen and Steinberg, 2018). In contrast, the degradative subdomain is marked by proteins which concentrate ubiquitinated membrane cargos for sorting to the lysosome including clathrin and the ESCRT complex (Cullen and Steinberg, 2018).

Underscoring the fundamental role of this cellular task, mutations in endosomal sorting proteins have been linked to a variety of human diseases (Maxfield, 2014; Kaur and Lakkaraju, 2018).

DNAJC13, and its *Caenorhabditis elegans* ortholog RME-8, is an endosomal protein that plays a critical role in this cargo sorting process (Zhang *et al*., 2001; Chang *et al*., 2004; Girard *et al*., 2005; Fujibayashi *et al*., 2008). DNAJC13 is the only known endosomal protein containing a DnaJ domain, and its interaction with the constitutively expressed heat shock protein 70 (HSC70; a member of the Hsp70 family) regulates the turnover of endosomal proteins which control sorting including SNX1 and clathrin (Chang *et al*., 2004; Girard *et al*., 2005; Popoff *et al*., 2009; Shi *et al*., 2009; Freeman *et al*., 2014). Consequently, loss of DNAJC13 results in missorting of both degrading and recycling cargos like the cation independent mannose-6- phosphate receptor, MIG-14/Wntless, Notch, the delta opioid receptor, and the beta-2 adrenergic receptor (Popoff *et al*., 2009; Shi *et al*., 2009; Gomez-Lamarca *et al*., 2015; Novy *et al*., 2024). DNAJC13 is also implicated in endosomal homeostasis, as loss of DNAJC13 causes aberrant enlargement of endosomes in human and *Drosophila melanogaster* cells, and loss of *C. elegans* RME-8 causes intermixing of normally spatially restricted endosomal subdomains governing recycling and degradation (Gomez-Lamarca *et al*., 2015; Norris *et al*., 2017; Novy *et al*., 2024). Consistent with a critical role in endosomal function, homozygous knockout of DNAJC13 in mice is embryonic lethal and heterozygous mice have decreased heart rate and hemoglobin (Groza *et al*., 2023). Additionally, point mutations in DNAJC13 have been linked to neurological diseases in humans including essential tremor and, potentially, Parkinson’s disease (Vilariño-Güell *et al*., 2014; Rajput *et al*., 2015; Deng *et al*., 2016; Deng and Siddique, 2017; Farrer *et al*., 2017).

Like other endosomal proteins, DNAJC13 must first localize to endosomes to function. Localization of DNAJC13 to endosomes is driven by its N-terminal Plekstrin Homology (PH)-like domain which can directly bind to the endosomal enriched phosphatidylinositol, phosphoinositol- 3-phosphate (PI(3)P) (Xhabija and Vacratsis, 2015). Deletion of the DNAJC13 N-terminus shifts its localization from endosomes to the cytoplasm, and point mutations within its N-terminal PH-like domain inhibit its localization in cells and block PI(3)P binding *in vitro* (Fujibayashi *et al*., 2008; Freeman *et al*., 2014; Xhabija and Vacratsis, 2015). Yet what regulates DNAJC13 localization to endosomes, and PI(3)P binding, is unknown. While DNAJC13 has been shown to bind other endosomal proteins including SNX1 and FAM21, these do not control its localization (Freeman *et al*., 2014; Xhabija and Vacratsis, 2015). One common mechanism that regulates endosomal proteins that bind directly to PI(3)P is that many have low affinity for PI(3)P as isolated monomers and have improved affinity for PI(3)P *in vitro,* and localization to endosomes in cells, when oligomerized (Klein *et al*., 1998; Hayakawa *et al*., 2004). This multivalency requirement for PI(3)P binding has been most clearly demonstrated for EEA1, where structural studies have defined a stalk region C-terminal to the FYVE domains that mediates homodimerization and positions the FYVE domains from two monomers such that each can simultaneously engage PI(3)P (Dumas *et al*., 2001). However, it is unknown if DNAJC13 has a multivalency requirement for PI(3)P-binding or if regions outside its N-terminus affect its ability to localize to endosomes.

Recent advances in structural modeling using AlphaFold (AF), and newer versions AF2 and AF3, have opened the door to creating specific, testable hypotheses about a protein’s structure-function relationships. We noted that the AF model of DNAJC13 predicted its C- terminal tail to be a 45 amino acid intrinsically disordered region (IDR) (**Figure 1A**) (Jumper *et al*., 2021; Varadi *et al*., 2022). As IDRs have a known role in protein regulation and autoinhibition, we hypothesized that this region may play a role in regulation of DNAJC13 function (Fenton *et al*., 2023). Thus, we set out to determine how localization of DNAJC13 to endosomes is regulated and how its distinct domains—including its N-terminal PH-like domain and its C-terminal tail—affect this localization.

**Figure 1.**
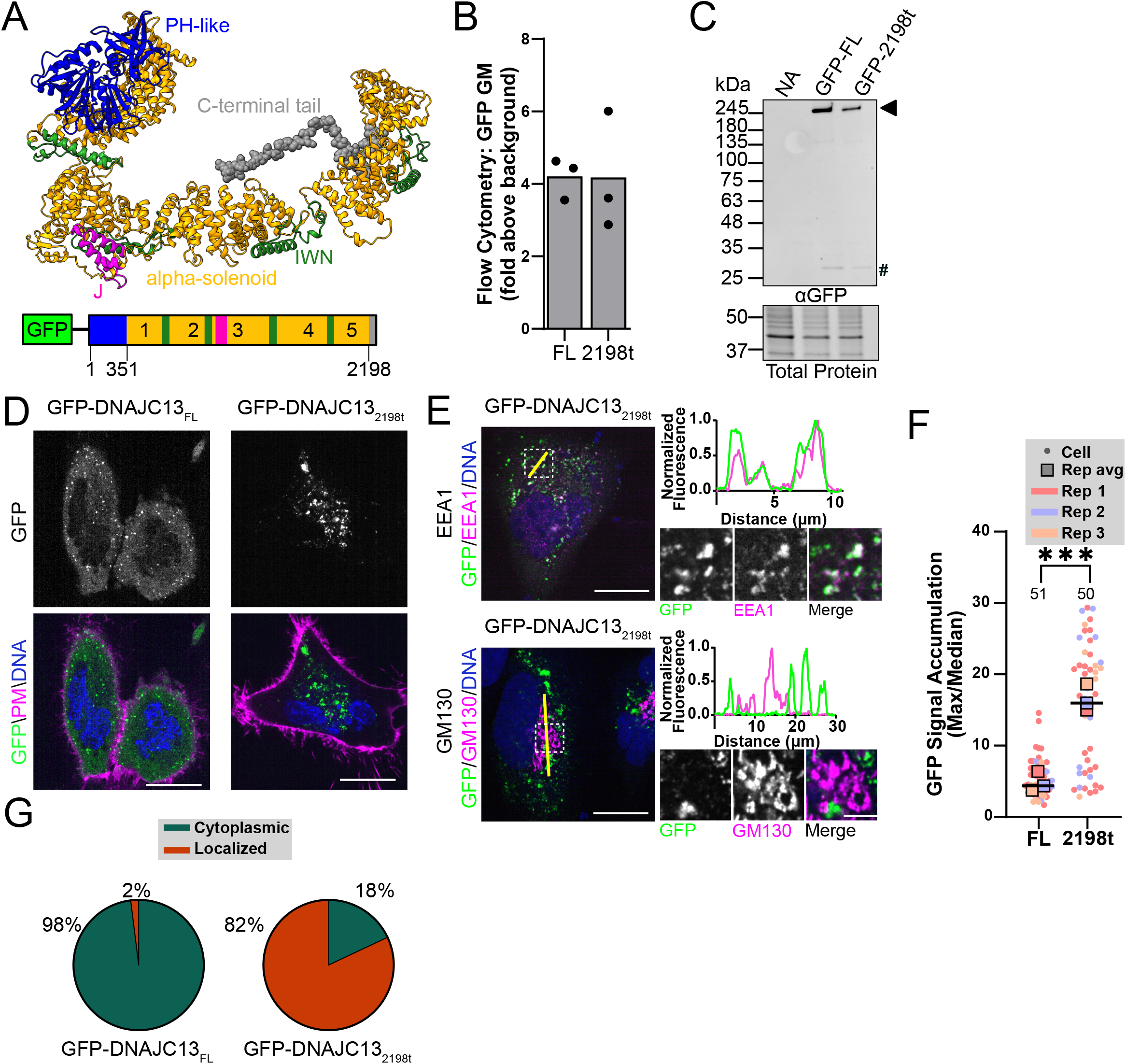
DNAJC13 disordered C-terminal tail controls its localization. ***A***, AlphaFold2.0 structure for human DNAJC13 (AF-O75165-F1-v4) (top) colored by domain (bottom), including the N-terminal PH-like domain (blue), five alpha solenoids (yellow) interspersed by repeating IWN motifs with potential regulatory function (Zhang *et al*., 2001; Norris *et al*., 2022) (dark green), a J domain (magenta) and C-terminal tail (grey, space filled residues). ***B***, Flow cytometry-based expression analysis of GFP-DNAJC13 constructs transfected into HeLa cells, assessed by geometric mean of GFP channel, displayed as fold above background signal from untransfected cells (n=3 biological replicates). ***C***, Representative western blot of transient expression of GFP-DNAJC13 constructs in HeLa cells, with a nontransfected control, with anti- GFP immunoblot (top) and total protein loading control (bottom), (n=3 biological replicates). The arrowhead marks GFP-DNAJC13 and the # marks free GFP. ***D***, Live spinning disk confocal microscopy of GFP-DNAJC13 constructs in HeLa cells. Imaged with CellMask plasma membrane stain (magenta) and Hoechst DNA stain (blue) (scale bar = 20 µm) (representative example from n=3 biological replicates). ***E***, Fixed immunofluorescent microscopy image of GFP- DNAJC13_2198t_ expressed in HeLa cells. Imaged with anti-GFP (Green), DAPI DNA stain (blue), and endosomal marker anti-EEA1 (magenta, top) or Golgi marker GM130 (magenta, bottom). Insets shown to the right (scale bar = 20 µm, 5 µm in inset), (representative example from n=3 biological replicates). Line-scans (yellow line) showing normalized fluorescent intensity of GFP (green) and EEA1 (magenta) or GM130 (magenta) signal are plotted along the line (right). ***F***, SuperPlot of cellular GFP signal accumulation metric (maximal GFP signal divided by median GFP signal) of individual cells with single cell data shown in circles and biological replicate averages plotted in squares (Lord *et al*., 2020). Total number of cells assessed is noted above the dataset (n=3 biological replicates, unpaired two-tailed t-test comparing biological replicate averages, p=0.0010). ***G***, Blinded analysis of live cell microscopy images of cells expressing DNAJC13_FL_ and DNAJC13_2198t_ for phenotype either being largely cytoplasmic (green) or localized to vesicles (orange). Cells scored are the same cells as those plotted in F.

## Results

### DNAJC13 disordered C-terminal tail controls its localization

We noted that the AF2 model of human DNAJC13 predicted its C-terminal tail, consisting of its final 45 amino acids, to be an IDR (**Figure 1A**). We next examined two other structural prediction programs, the disorder predictor JRonn and five additional AF3 models, which also predicted the C-terminal tail of DNAJC13 to be disordered (**Figure S1A**) (Waterhouse *et al*., 2009; Troshin *et al*., 2011; Abramson *et al*., 2024). As IDRs commonly serve regulatory functions, we hypothesized that the C-terminal tail of DNAJC13 could affect its localization to endosomes (Fenton *et al*., 2023).

To test this hypothesis, we designed several DNAJC13 constructs using the same N-terminal GFP tagging scheme as those used in the literature (Fujibayashi *et al*., 2008; Xhabija *et al*., 2011; Freeman *et al*., 2014; Yoshida *et al*., 2018): full-length GFP-DNAJC13 (DNAJC13_FL_) or GFP-DNAJC13 lacking its 45 amino acid C-terminal tail (DNAJC13_2198t_). We first analyzed the relative expression of these constructs by flow cytometry and found they express at similar levels (**Figure 1B**). Additionally, by western blot we saw minimal evidence of proteolysis and liberation of free GFP (**Figures 1C**, **S1B**). We also examined full-length DNAJC13 with a C-terminal GFP (DNAJC13_FL_-GFP) but found that it expressed poorly (less than 10% of the expression of DNAJC13 with an N-terminal GFP tag), which did not allow for further analysis (**Figure S1C**).

We then sought to determine the localization of these GFP-DNAJC13 constructs in cells using live microscopy and found, similar to previous observations, that overexpressed DNAJC13_FL_ localized to both the cytoplasm and endosomes (**Figures 1D**, **S1D**) (Fujibayashi *et al*., 2008; Freeman *et al*., 2014). Strikingly, DNAJC13_2198t_ was highly localized to vesicles with minimal cytoplasmic background (**Figure 1D**). As DNAJC13/RME-8 localizes to early endosomes, we turned to immunofluorescent microscopy to determine the identity of the DNAJC13-positive structures (Zhang *et al*., 2001; Girard *et al*., 2005; Fujibayashi *et al*., 2008; Shi *et al*., 2009; Xhabija and Vacratsis, 2015; Novy *et al*., 2024). Using the early endosomal marker EEA1 and the Golgi marker GM130, we confirmed that GFP-DNAJC13-positive vesicles are indeed early endosomes (**Figures 1E**, **S1D**). Thus, by both live and fixed imaging, we found that removal of the DNAJC13 C-terminal tail enhanced its localization to endosomes.

To further characterize the enhanced vesicular localization in GFP-DNAJC13_2198t_, we performed two orthogonal methods of analysis. First, to quantitatively score cells, we devised a GFP signal accumulation metric where each cell’s maximal fluorescence is divided by its median fluorescence. In this metric a score of 1 would indicate the signal is homogeneous throughout the cell, much like free GFP, while a high score indicates a localized protein.

Second, we performed blinded qualitative analysis to assess GFP signal in cells as either “cytoplasmic,” containing highly cytoplasmic GFP and few distinct GFP-positive vesicles, or “localized,” containing GFP predominantly localized to vesicles with little to no cytoplasmic GFP.

Using the quantitative GFP accumulation metric, we found that DNAJC13_2198t_ had ∼3.6- fold higher score than DNAJC13_FL_ (**Figure 1F**). Blinded qualitative analysis confirmed these findings, with only 2% of cells expressing DNAJC13_FL_ showing a predominant vesicular localization compared to 82% of cells expressing DNAJC13_2198t_ (**Figure 1G**). The orthogonal nature of these methods also allowed for direct comparison between the qualitative (blinded scoring) and quantitative (signal accumulation metric) analyses, which showed broad agreement between our two approaches, with cells showing localized GFP signal having a higher signal accumulation score (**Figure S1E**). Together, these data demonstrate that DNAJC13 localization to early endosomes is negatively regulated by its disordered C-terminal tail.

### YLT residues in C-terminal tail control endosomal localization

We next asked which part of the DNAJC13 C-terminal tail was necessary to control its localization to endosomes. To narrow down the scope of our search, we first assessed the evolutionary conservation of the last 45 amino acids of DNAJC13—those predicted by AF2 and AF3 to be disordered—by calculating a relative conservation score using the Ensembl database of vertebrate orthologues (plus *C. elegans and D. melanogaster*) (Waterhouse *et al*., 2009; Harrison *et al*., 2024). We found that the first half of the tail was more highly conserved than the second half (**Figure 2A**). Consequently, we focused on this conserved region and used alanine scanning to mutate blocks of three residues at a time to probe for which amino acids were important in controlling DNAJC13 localization (**Figure 2A**, brackets). Analysis of these constructs showed they were expressed at similar levels without significant proteolysis (**Figures S2A-B**).

**Figure 2.**
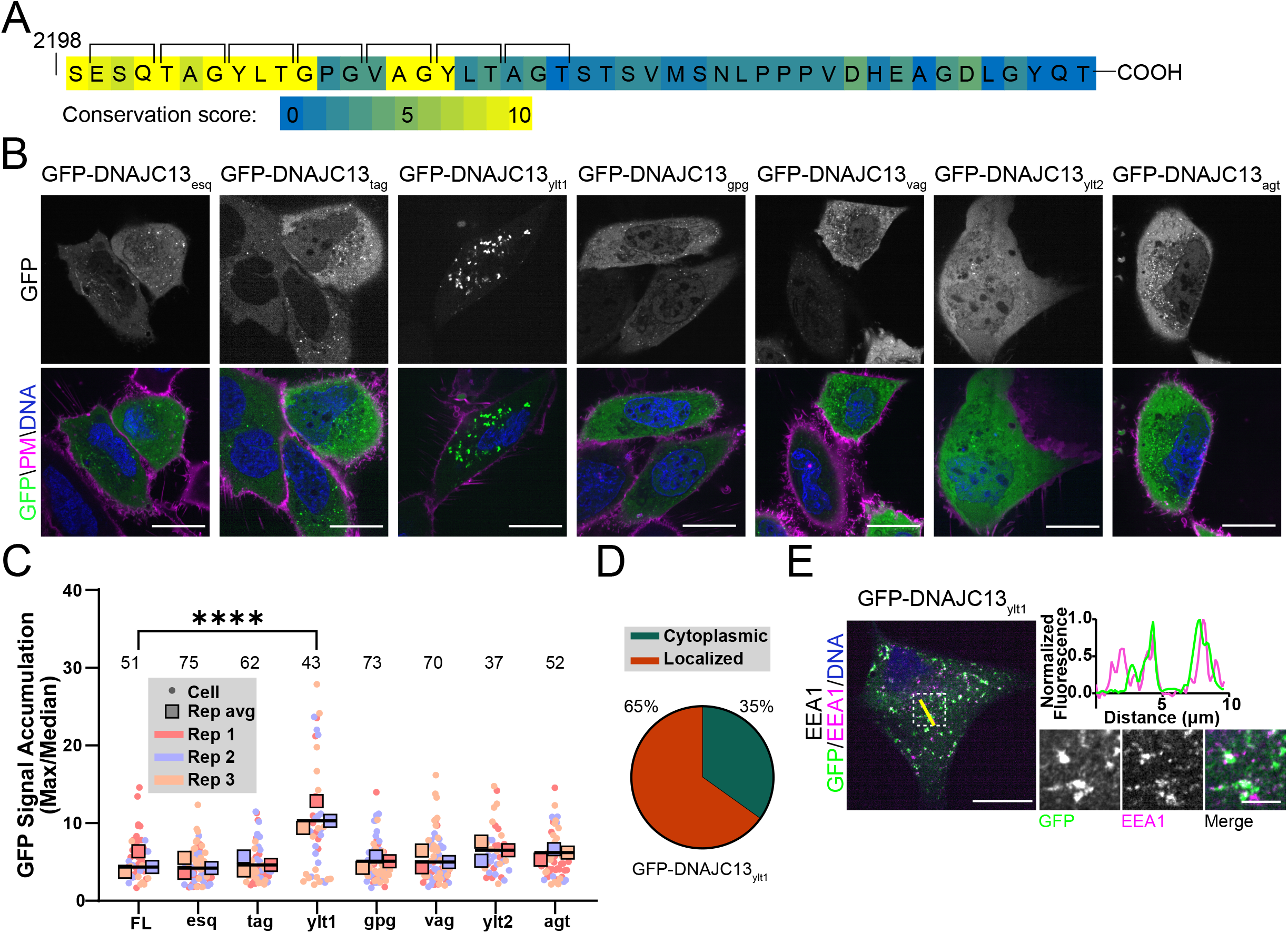
YLT residues in C-terminal tail control endosomal localization. ***A***, Relative conservation analysis of the DNAJC13 C-terminal tail (45 residues) amongst all orthologues in Ensemble vertebrate (plus *C. elegans and D. melanogaster*) database (less conserved = more blue; more conserved = more yellow). Brackets above indicate regions for triplet alanine scanning. ***B***, Live spinning disk confocal microscopy of triplet scan mutagenesis, expressed in HeLa cells. Imaged with CellMask plasma membrane stain (magenta) and Hoechst DNA stain (blue) (scale bar = 20 µm) (representative example from n=3 biological replicates). ***C***, SuperPlot of cellular GFP signal accumulation metric of individual cells with single cell data shown in circles and biological replicate averages plotted in squares. Total number of cells assessed is noted above the dataset (n=3 biological replicates, one-way unpaired ANOVA comparing biological replicate averages with Dunnett’s multiple comparisons corrections, all vs DNAJC13_FL_, p<0.0001 for DNAJC13_ylt1_, ns for all other mutants). ***D***, Blinded analysis of live cell microscopy images of cells expressing DNAJC13_ylt1_ for phenotype either being largely cytoplasmic (green) or localized to vesicles (orange). Cells scored are the same cells as those plotted in C. ***E***, Fixed immunofluorescent microscopy image of GFP-DNAJC13_ylt1_ expressed in HeLa cells. Imaged with anti-GFP (green), endosomal marker anti-EEA1 (magenta), and DAPI DNA stain (blue) with insets shown to the right (scale bar = 20 µm, 5 µm in inset), (representative example from n=3 biological replicates). A line-scan (yellow line) showing normalized fluorescent intensity of GFP (green) and EEA1 (magenta) signal are plotted along the line (right).

Using live cell microscopy and the quantitative GFP signal accumulation metric, we assessed these constructs for localization and found that only one mutant, DNAJC13_ylt1_ (Y2206A, L2207A, T2208A) significantly increased vesicular accumulation (∼2.4-fold above DNAJC13_FL_; **Figures 2B-C**). Consistent with this observation, blinded qualitative analysis of DNAJC13_ylt1_ found 65% of the cells contained GFP-DNAJC13 signal localized to predominantly vesicles (**Figure 2D**). We again confirmed endosomal localization of DNAJC13_ylt1_ with immunofluorescence imaging using EEA1 and GM130 probes (**Figures 2E**, **S2C**). We noted that both scoring metrics showed the DNAJC13_ylt1_ phenotype was less penetrant than DNAJC13_2198t_ (signal accumulation: 10.30 vs 15.98, respectively; localization phenotype: 65% vs 82%, respectively). Closer examination of the DNAJC13 C-terminus revealed a second instance of the YLT sequence (Y2215, L2216, T2217), called DNAJC13_ylt2_, downstream of DNAJC13_ylt1_. Quantitative analysis of DNAJC13_ylt2_ localization showed a non-significant trend toward enhanced vesicular localization (∼1.5-fold enhancement). Thus, it is possible that the difference in effect size between DNAJC13_2198t_ and DNAJC13_ylt1_ could be explained by the minor contribution of DNAJC13_ylt2_ in control of DNAJC13 localization. Lastly, we examined if YLT1 from human DNAJC13 was conserved in commonly used model systems, *C. elegans* and *D. melanogaster*, and found that the motif is intact within the *D. melanogaster* homologue but only partially present in the *C. elegans* homologue (**Figure S2D**). Together, our data suggests a model in which the C-terminal tail, driven primarily by a YLT sequence (Human: 2206-2208) regulates DNAJC13’s endosomal localization.

### J domain co-regulates DNAJC13 localization

To gain insight into what restricts DNAJC13 localization to the cytoplasm, we next sought to identify the protein-protein interactions of DNAJC13_FL_. To this end, we separately purified free GFP or GFP-DNAJC13_FL_ using an anti-GFP nanobody and performed quantitative proteomics with tandem mass tag (TMT) labeling (**Table S1**). In analyzing proteins specifically co-purified with DNAJC13_FL_, we found that many of the interactors were in the Hsp70 pathway— either part of the Hsp70 family (HSPA8/HSC70, HSPA1A/HSP70, HSPA9/GRP75) or Hsp70 co-chaperones (BAG2, STUB1/CHIP, and HSPA4) (**Figure 3A**, red and orange circles, respectively). These findings are consistent with previous observations showing that the J domain of DNAJC13 interacts with HSC70 (Chang *et al*., 2004; Girard *et al*., 2005; Ryu *et al*., 2020). Notably, we did not observe interactions with the DNAJC13 binding proteins FAM21 or SNX1, and we attribute this to the majority of GFP-DNAJC13_FL_ residing in the cytoplasm and thus likely not interacting with these endosomal proteins (Shi *et al*., 2009; Freeman *et al*., 2014).

**Figure 3.**
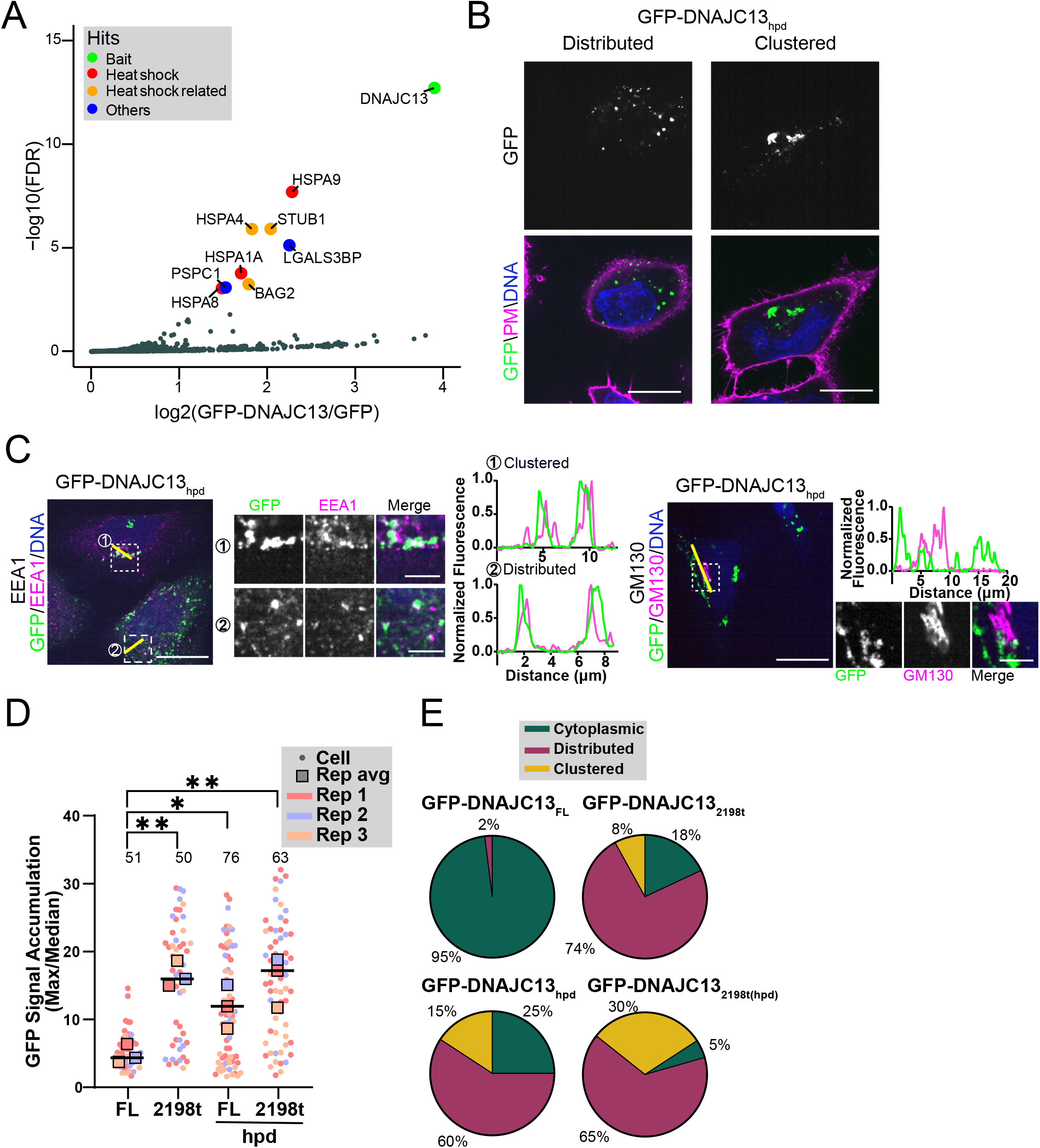
J domain co-regulates DNAJC13 localization. ***A***, Volcano plot of GFP-DNAJC13_FL_ proteomics, as compared to a GFP control (n=2 biological replicates). Hits are annotated as being bait protein (green), heat shock proteins (red), heat shock accessory proteins (orange), or other (blue). ***B***, Live cell spinning disk confocal microscopy images of GFP-DNAJC13_hpd_ in HeLa cells exhibiting distributed (left) and clustered (right) endosomes. Imaged with CellMask plasma membrane stain (magenta) and Hoechst DNA stain (blue) (scale bar = 20 µm) (phenotypic representative examples from n=3 biological replicates). ***C***, Fixed immunofluorescent microscopy image of GFP-DNAJC13_hpd_ expressed in HeLa cells. Imaged with anti-GFP (green), DAPI DNA stain (blue), and endosomal marker anti-EEA1 (magenta, left) or Golgi marker anti- GM130 (magenta, right). Insets shown to the right (scale bar = 20 µm, 5 µm in insets), (representative example from n=3 biological replicates). Line-scans (yellow lines) for each inset showing normalized fluorescent intensity of GFP (green) and EEA1 (magenta) or GM130 (magenta) signal are plotted along the lines (right). ***D***, SuperPlot of cellular GFP signal accumulation metric of individual cells with single cell data shown in circles and biological replicate averages plotted in squares. Total number of cells assessed is noted above the dataset (n=3 biological replicates, one-way unpaired ANOVA comparing biological replicate averages with Dunnett’s multiple comparisons corrections, all to DNAJC13_FL_, p=0.0020 (DNAJC13_2198t_), 0.0.0322 (DNAJC13hpd), 0.0028 (DNAJC13_2198t(hpd)_)). ***E***, Blinded analysis of live cell microscopy images of cells expressing DNAJC13_FL_, DNAJC13_2198t_, DNAJC13hpd, and DNAJC13_2198t(hpd)_ for phenotype being either: largely cytoplasmic (green), localized to distributed endosomes (purple), or localized to endosomes clustered to a perinuclear region (yellow). Cells scored are the same cells as those plotted in D.

As the majority of the DNAJC13 binding proteins were HSC70 pathway proteins, we reasoned that the activity of the J domain of DNAJC13 could be critical to the cytoplasmic localization of GFP-DNAJC13_FL_. To test this hypothesis, we created constructs in which the HPD residues in the J domain, which are critical for binding HSC70 and stimulating HSC70 ATPase activity, were mutated to alanines (termed DNAJC13hpd and a dual mutant, DNAJC13_2198t(hpd)_) (Chamberlain and Burgoyne, 1997; Morgan *et al*., 2001; Yan *et al*., 2002; Tummala *et al*., 2016). These constructs expressed at similar levels with minimal proteolysis (**Figures S3A-B**).

Similar to DNAJC13_2198t_, both DNAJC13hpd and DNAJC13_2198t(hpd)_ showed strong localization to endosomes with little DNAJC13 residing in the cytoplasm (**Figures 3B-C**, **S3C-D**). The observation that loss of J domain function increased DNAJC13 localization to vesicles was supported by the GFP signal accumulation metric (DNAJC13hpd ∼2.7-fold above DNAJC13_FL_, DNAJC13_2198t(hpd)_ ∼3.9-fold above DNAJC13_FL_; **Figure 3D**). Interestingly, we observed that in a subset of the DNAJC13hpd and DNAJC13_2198t(hpd)_ expressing cells, the GFP- DNAJC13-positive endosomes clustered in a perinuclear region that was distinct from the Golgi (**Figures 3B-C**, **S3C-D**). A similar phenotype of endosomal clustering has also been observed upon manipulation of proteins which functionally interact with DNAJC13—the WASH complex and clathrin (Bennett *et al*., 2001; Gomez *et al*., 2012).

To further assess endosomal redistribution, we performed blinded analysis in which cells were scored for GFP signal as being predominantly cytoplasmic, localized to distributed vesicles, or localized to clustered vesicles. We found no instances of the endosomal clustering phenotype in cells expressing GFP-DNAJC13_FL_, while cells expressing DNAJC13_2198t_ or DNAJC13hpd showed similar proportions of distributed and clustered endosomal vesicles (DNAJC13_2198t_: 74% distributed, 8% clustered; DNAJC13hpd: 60% distributed, 15% clustered; **Figure 3E**). Consistent with an additive effect of these two mutations, we found a larger percentage of cells (30%) expressing GFP-positive DNAJC13_2198t(hpd)_ showed an endosomal clustered phenotype (**Figure 3E**). Cross-comparison of the two metrics show there is no correlation between endosomal clustering and signal accumulation (**Figure S3E**). These observations suggest that there are two control points for DNAJC13 localization to endosomes: YLT motif(s) in its C-terminal tail and its J domain. Additionally, our observations suggest that similar to disruption of WASH or clathrin function, overexpression of DNAJC13 carrying these activating mutations can act in a dominant negative manner to affect endosomal distribution in the cell (Bennett *et al*., 2001; Gomez *et al*., 2012).

### C-terminal tail and J domain act through PH-like domain to enhance PI(3)P binding

We then sought to analyze the mechanism by which the J domain and C-terminal mutants enhance DNAJC13 localization to endosomes. DNAJC13 is known to localize to endosomes through a single PH-like domain in its N-terminus (first ∼100 residues) (Xhabija *et al*., 2011; Xhabija and Vacratsis, 2015). Thus, we considered the possibility that the J domain and C-terminal IDR were modulating the ability of the N-terminal PH-like domain to bind to PI(3)P.

To test this, we examined binding of DNAJC13 in detergent lysates to agarose beads conjugated to phosphoinositides. As had been observed previously, we found that DNAJC13_FL_ bound efficiently to PI(3)P and did not bind to the negative control, phosphatidylinositol phosphate (PIP) (**Figures 4A**, **S4A**) (Xhabija *et al*., 2011; Xhabija and Vacratsis, 2015). We then examined the DNAJC13 mutations that enhanced endosomal localization (DNAJC13_2198t_, DNAJC13_ylt1_, and DNAJC13hpd) and found increased binding compared to DNAJC13_FL_ (**Figures 4A, S4A**). Quantification of this result showed that DNAJC13_2198t_ and DNAJC13hpd bound PI(3)P decorated resins ∼five-fold better than DNAJC13_FL_ (**Figure 4B**). Comparatively, we observed an ∼three-fold better PI(3)P binding of DNAJC13_ylt1_ compared to DNAJC13_FL_, although this did not reach statistical significance (**Figure 4B**). DNAJC13_ylt1_ had lower scores in both quantitative and qualitative analysis of its localization in cells compared to the other mutants, which is consistent with its weaker PI(3)P binding *in vitro*.

**Figure 4.**
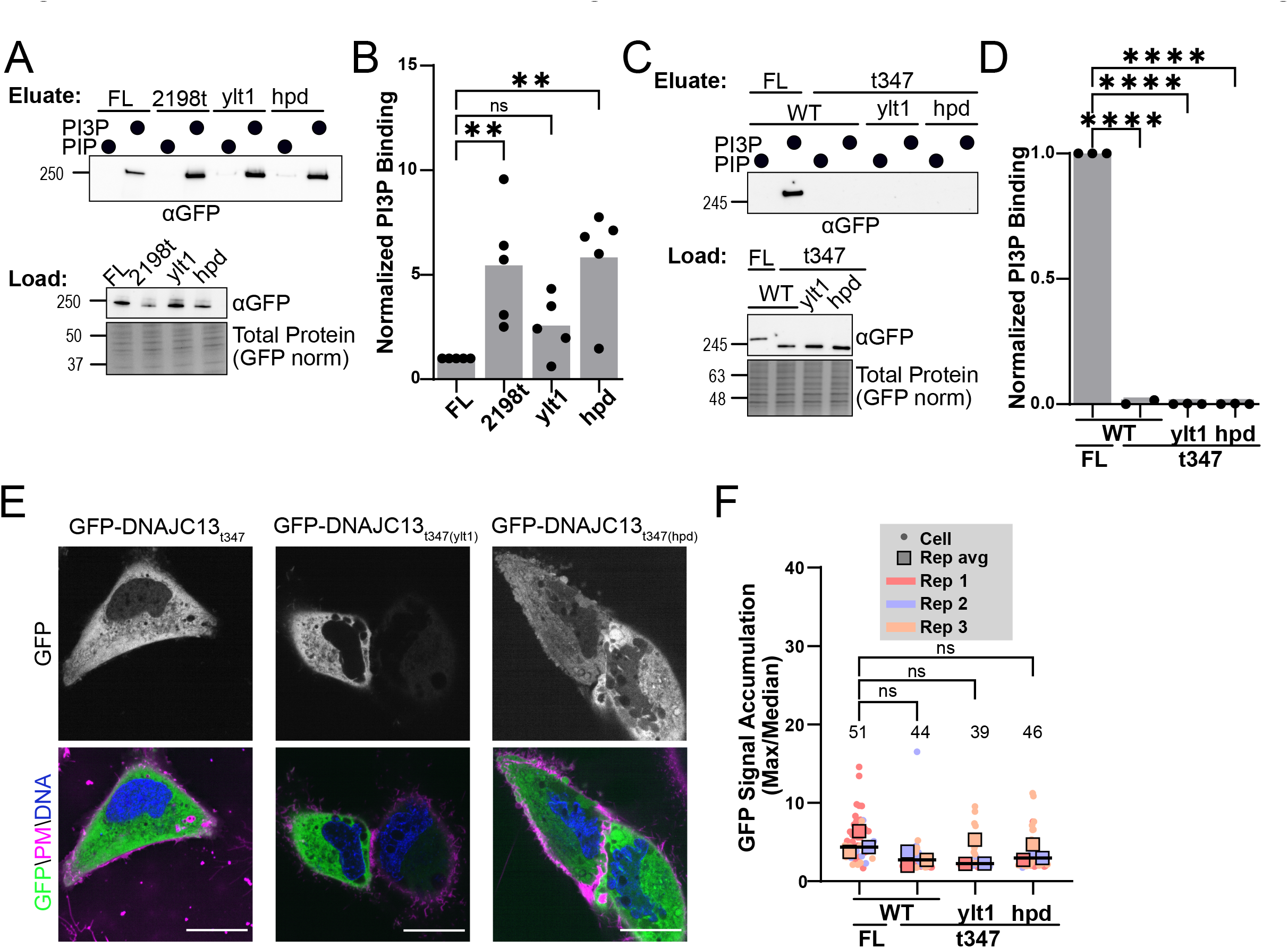
C-terminal tail and J domain act through PH-like domain to enhance PI(3)P binding. ***A***, Western blots of PIP resin eluates for DNAJC13_FL_ and activating mutants. GFP- DNAJC13_FL_, GFP-DNAJC13_2198t_, GFP-DNAJC13_ylt1_, and GFP-DNAJC13_hpd_ were expressed in HEK293 cells and lysates, normalized by flow cytometry for GFP expression, and were bound to PIP (control) and PI(3)P decorated agarose resins. Loads and eluates were run on SDS-PAGE (load total protein stain, bottom) and immunoblotted for anti-GFP (load, middle; eluate, top). ***B***, Quantification of PI(3)P pulldowns in A, normalized to load and the full-length pulldown (n=4 biological replicates, one-way unpaired ANOVA with Dunnett’s multiple comparisons corrections, all vs DNAJC13_FL_, p= 0.0085 (DNAJC13_2198t_), 0.4915 (DNAJC13_ylt1_), 0.0046 (DNAJC13hpd)). ***C***, Western blots of PIP resin eluates of DNAJC13 lacking PH-like domains. GFP-DNAJC13_FL_, GFP-DNAJC13_t347_, GFP-DNAJC13_t347(ylt1)_, and GFP-DNAJC13_t347(hpd)_ were expressed in HEK293 cells and lysates, normalized by flow cytometry for GFP expression, were bound to PIP (control) and PI(3)P decorated agarose resins. Loads and eluates were run on SDS-PAGE (load total protein stain, bottom) and immunoblotted for anti-GFP (load, middle; eluate, top). ***D***, Quantification of PI(3)P pulldowns in C, normalized to load and the full-length pulldown (n=3 biological replicates for all but DNAJC13_t347_ which has n=2, one-way unpaired ANOVA with Dunnett’s multiple comparisons corrections vs DNAJC13_FL_, p<0.0001 for all comparisons). ***E***, Live cell spinning disk confocal microscopy of GFP-DNAJC13_t347_, GFP- DNAJC13t34(ylt1), GFP-DNAJC13_t347(hpd)_ in HeLa cells. Imaged with CellMask plasma membrane stain (magenta) and Hoechst DNA stain (blue) (scale bar = 20 µm), (representative example from n=3 biological replicates). ***F***, SuperPlot of cellular GFP signal accumulation metric of individual cells with single cell data shown in circles and biological replicate averages plotted in squares. Total number of cells assessed is noted above the dataset (n=3 biological replicates, one-way unpaired ANOVA comparing biological replicate averages with Dunnett’s multiple comparisons corrections, all vs DNAJC13_FL_, ns for all).

We next tested if the enhanced PI(3)P binding we observed upon mutation of the J domain or C-terminal tail required the N-terminal PH-like domain. A recent AF2 analysis of the *C. elegans* homologue RME-8 identified that the first 300 amino acids contain not one but three folds which each resemble PH-like domains (Norris *et al*., 2022). While it is not known if these second and third PH-like domains bind PI(3)P—and it is notable that single point mutation in the first PH-like domain of DNAJC13 fully blocked PI(3)P binding *in vitro* and endosomal localization in cells (Xhabija and Vacratsis, 2015)—we decided to examine constructs lacking all three predicted PH-like domains in the wild type and mutated contexts (truncation of residues 1-347, termed DNAJC13_t347_, DNAJC13_t347(ylt1)_ and DNAJC13_t347(hpd)_).

These constructs expressed at similar levels with minimal proteolysis (**Figures S4B-C**). We found that removal of the DNAJC13 N-terminus (DNAJC13_t347_) blocked binding of DNAJC13 to PI(3)P *in vitro* and localization to endosomes in cells, and that the J domain and C-terminal mutants did not rescue these phenotypes (**Figures 4C-D**). These findings were validated by the signal accumulation metric which showed a trend toward a lower score in cells expressing DNAJC13_t347_ constructs compared to cells expressing DNAJC13_FL_ and thus support the model for an absolute requirement of the DNAJC13 PH-like domain for its localization (**Figures 4E-F**). Together, these results demonstrate that the DNAJC13 C-terminal tail and J domain act in the same pathway as its N-terminal PH-like domain to control DNAJC13 binding to PI(3)P *in vitro* and localize to endosomes in cells.

### PH-like domain requires oligomerization for efficient PI(3)P binding and endosomal localization

We next considered a possible mechanism by which relatively distal parts of the DNAJC13 protein could affect the function of its N-terminal PH-like domain. One of the known regulatory mechanisms for some proteins that bind PI(3)P is a requirement for multivalency. For example, the FYVE domains of EEA1, Hrs, and Frabin localize to endosomes poorly as isolated domains but localize efficiently when artificially oligomerized (Hayakawa *et al*., 2004). For EEA1, structural studies have shown a stalk region upstream of the FYVE domain mediate dimerization between two monomers to position tandem FYVE domains for PI(3)P binding (Dumas *et al*., 2001). Additionally, recent studies of the *C. elegans* homolog RME-8 have proposed a model in which oligomerization of RME-8 is a critical part of its endosomal catalytic cycle (Norris *et al*., 2022). Thus, we wanted to determine if the PH-like domain of DNAJC13 was sufficient in isolation to localize to endosomes and bind PI(3)P or if it, like a subset of other endosomal proteins, required oligomerization.

We designed constructs to express the N-terminal DNAJC13_PH-like_ in isolation (1-351, termed DNAJC13_351t_) and additionally made constructs fusing the PH-like domain to established dimerization and tetramerization motifs (DNAJC13_351t_-dimer and DNAJC13_351t_-tetramer, respectively) (**Figure 5A**) (Khairil Anuar *et al*., 2019). We first analyzed the binding of these constructs to PI(3)P beads in detergent lysate. Interestingly, we were unable to detect appreciable binding of the isolated PH-like domain to PI(3)P beads (**Figures 5B-C**, **S5A**).

**Figure 5.**
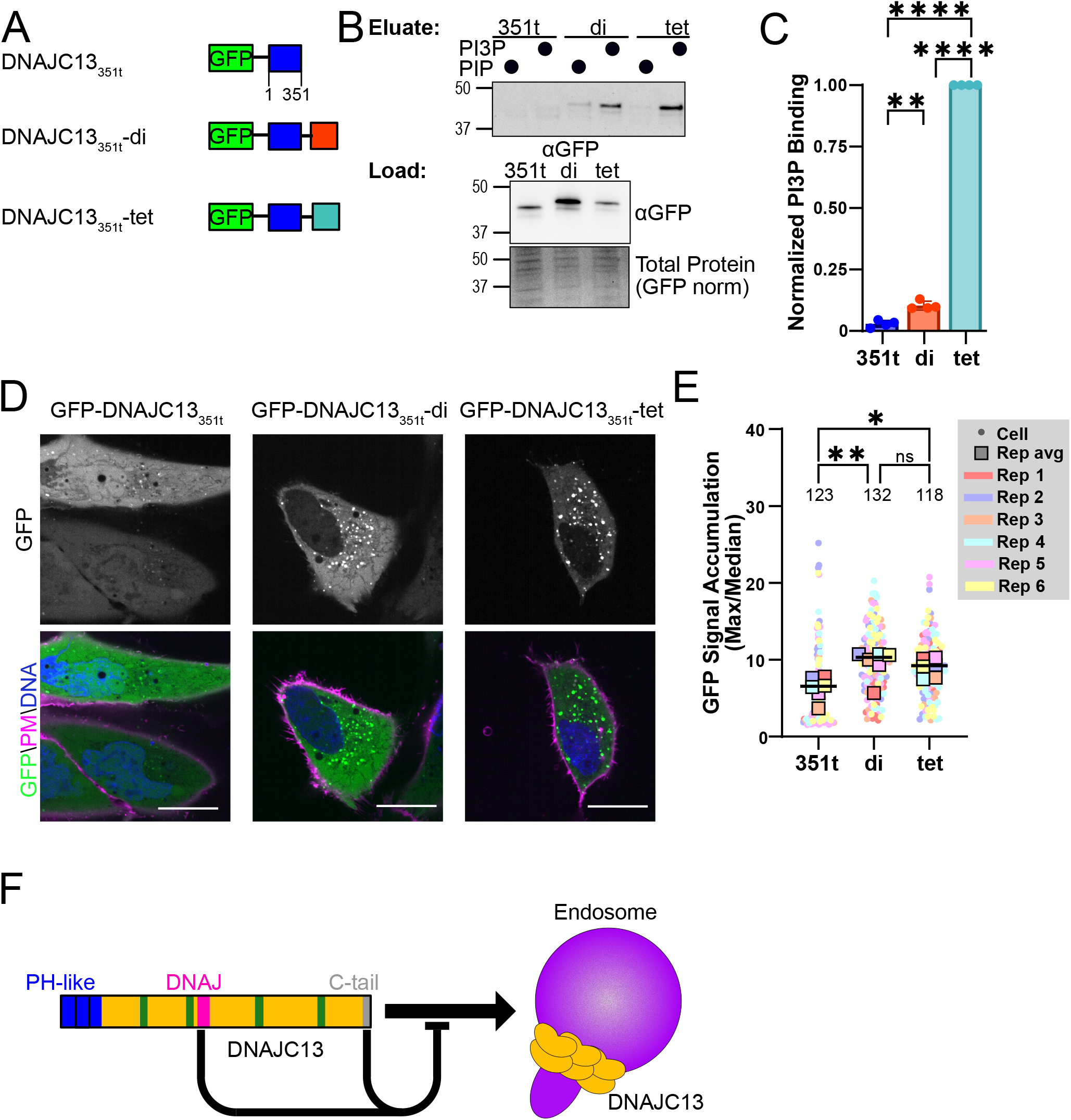
PH-like domain requires oligomerization for efficient PI(3)P binding and endosomal localization. ***A***, Domain schematics of GFP-tagged constructs containing only the PH-like domains (DNAJC13_351t_) and constructs containing exogenous dimerization (DNAJC13_351t_-dimer) and tetramerization (DNAJC13_351t_-tetramer) motifs. ***B***, Western blots of PIP resin eluates for DNAJC13_351t_ constructs. GFP-DNAJC13_351t_ constructs were expressed in HEK293 cells and lysates, normalized by flow cytometry for GFP expression, and were bound to PIP (control) and PI(3)P decorated agarose resins. Loads and eluates were run on SDS-PAGE (load total protein stain, bottom) and immunoblotted for anti-GFP (load, middle; eluate, top). ***C***, Quantification of PI(3)P pulldowns in B, normalized to load and the DNAJC13_351t_-tetramer pulldown (n=4 biological replicates, one-way paired ANOVA with Tukey’s multiple comparisons corrections, p = 0.0025 (DNAJC13_351t_ vs DNAJC13_351t_-dimer), <0.0001 (DNAJC13_351t_ vs DNAJC13_351t_-tetramer), <0.0001 (DNAJC13_351t_-dimer vs DNAJC13_351t_-tetramer)). ***D***, Live cell spinning disk confocal microscopy of GFP-DNAJC13_351t_ constructs in HeLa cells. Imaged with CellMask plasma membrane stain (magenta) and Hoechst DNA stain (blue) (scale bar = 20 µm), (representative example from n=3 biological replicates). ***E***, SuperPlot of cellular GFP signal accumulation metric of individual cells with single cell data shown in circles and biological replicate averages plotted in squares. Total number of cells assessed is noted above the dataset (n=6 biological replicates, one-way unpaired ANOVA comparing biological replicate averages with Tukey’s multiple comparisons corrections, p= 0.0087 (DNAJC13_351t_ vs DNAJC13_351t_-dimer), 0.0258 (DNAJC13_351t_ vs DNAJC13_351t_-tetramer), 0.85 (DNAJC13_351t_-dimer vs DNAJC13_351t_-tetramer). ***F***, Cartoon schematic of proposed mechanism whereby DNAJC13’s J domain and YLT motif in the C-terminal tail inhibit oligomerization and localization to endosomes.

However, binding increased when the DNAJC13_PH-like_ was dimerized, and was even further enhanced with tetramerization (**Figures 5B-C**, **S5A**). These observations demonstrate that similar to other PI(3)P-binding proteins, the DNAJC13_PH-like_ domain binds weakly to PI(3)P as a monomer and its binding is enhanced upon oligomerization.

To investigate the localization of the DNAJC13_PH-like_ domain in cells, we first confirmed the isolated, dimeric, and tetrameric constructs expressed at similar levels, slightly higher than the full-length construct, with minimal proteolysis (**Figures S5B-C**). By live cell microscopy, DNAJC13_PH-like_ looked similar to DNAJC13_FL_, with the GFP signal largely cytoplasmic with some vesicular localization (**Figures 5D**). Consistent with our *in vitro* assays, the dimerization or tetramerization of the DNAJC13_PH-like_ enhanced its localization to vesicles which were confirmed to be endosomes with immunofluorescent imaging (**Figures 5D**, **S5D**). Using the GFP signal accumulation metric, we confirmed that DNAJC13_PH-like_-dimer and DNAJC13_PH-like_-tetramer localized to vesicles more strongly than the isolated DNAJC13_PH-like_ (**Figure 5E**). We did not observe a difference in the degree of localization between the dimeric and tetrameric constructs, potentially due to saturation of PI(3)P binding sites in cells. Additionally, we observed no signs of endosomal clustering with these constructs, suggesting that while the PH-like domain controls localization, other parts of DNAJC13 affect its function in cells. Together, these data demonstrate that similar to other PI(3)P binding proteins, the DNAJC13 PH-like domain binds weakly to PI(3)P in isolation and its binding to PI(3)P—and therefore ability to localize to endosomes—can be enhanced by oligomerization.

## Discussion

Our findings demonstrate that DNAJC13 localization in cells is controlled by the cumulative function of three different domains: its N-terminal PH-like domain, which weakly binds PI(3)P, as well as its J domain and C-terminus, which act functionally upstream of the PH-like domain to oppose DNAJC13 localization to endosomes. Furthermore, we show that the poor endosomal localization of the DNAJC13 PH-like domain to endosomes can be improved by oligomerization, an observation consistent with a subset of other PI(3)P binding domains as well as recent findings that suggest the *C. elegans* homologue, RME-8, oligomerizes as part of its functional lifecycle (Klein *et al*., 1998; Dumas *et al*., 2001; Hayakawa *et al*., 2004; Norris *et al*., 2022). Thus, in a working model we propose that DNAJC13 exists in an equilibrium between a cytoplasmic inhibited state and an oligomeric state that can localize efficiently to endosomes, with the transition between these states being controlled by a YLT motif in the C-terminal tail and the catalytic triad, HPD, in the J domain (**Figure 5F**).

### PI(3)P Binding Domains and Oligomerization

Our data demonstrate that the isolated PH-like domain of DNAJC13 localizes poorly to endosomes in cells and weakly to PI(3)P *in vitro*, and this can be partially rescued through artificial oligomerization. This observation parallels what has been found for other PI(3)P binding domains like that from HRS, EEA1 and Frabin (Dumas *et al*., 2001; Hayakawa *et al*., 2004). For example, the PI(3)P binding domain in HRS associates with endosomes poorly as an isolated monomer but efficiently when artificially dimerized (Hayakawa *et al*., 2004). Multivalency in phosphatidylinositol binding is not limited to FYVE domains as a similar requirement has been shown for the PH-domain in dynamin (Klein *et al*., 1998; Lemmon, 2007). While not all PI(3)P binding proteins require oligomerization to bind to PI(3)P and endosomes (e.g., WDFY1 and endofin), multivalency—such as with EEA1—has been shown to allow for another layer of regulation (Blatner *et al*., 2004; Kim *et al*., 2005; Ramanathan and Ye, 2012). While oligomerization can assist PI(3)P binding in some cases, other proteins like DFCP1/ZFYVE1 have naturally occurring tandem FYVE domains that are both required for high affinity PI(3)P binding (Cheung *et al*., 2001; Hayakawa *et al*., 2004). In this light it is interesting to note that a recent AF analysis of *C. elegans* RME-8 revealed the presence of three tandem PH-like domains in the first ∼300 residues (Norris *et al*., 2022). While future studies will be required to determine if these domains provide multivalency in PI(3)P binding, previous studies showed that a single point mutation in first PH-like domain was sufficient to block PI(3)P binding and our study showed all three PH-like domains (1-351) bound weakly to PI(3)P *in vitro* and endosomes in cells (Xhabija and Vacratsis, 2015). Together, our study demonstrates that similar to other PI(3)P binding domains, the PI(3)P binding domain in DNAJC13 operates poorly in isolation and is enhanced by oligomerization (Dumas *et al*., 2001; Hayakawa *et al*., 2004).

The nature of our experiments allows for direct comparison of PI(3)P binding between the DNAJC13 PH-like domain in isolation, dimerized, and tetramerized, or in wild-type and mutationally activated, full-length DNAJC13 constructs. One observation that arose from these comparisons is that full-length constructs bound to PI(3)P resins much better than the tetramerized PH-like domain (**Figure S5A**). One potential explanation for this finding is that another PI(3)P binding protein functions cooperatively with DNAJC13 in binding PI(3)P resins; however, we consider this unlikely given the absence of such a protein in our proteomics results. If PI(3)P binding was cooperative with another protein, the most likely candidate would be the DNAJC13 binding protein SNX1; however, SNX1 binds DNAJC13/RME-8 in its middle region (*C. elegans*: 1388-1950) and we show that loss of the N-terminal PH-like domains completely blocks DNAJC13 binding to PI(3)P *in vitro* and endosomes in cells, suggesting that this interaction by itself cannot localize DNAJC13 to endosomes (Shi *et al*., 2009). An alternate interpretation of our findings is that full-length DNAJC13 spontaneously forms larger order assemblies (>4-mer) *in vitro* which enhance PI(3)P binding through multivalency. Notably, GFP- DNAJC13_FL_ and GFP-DNAJC13_PH-like_-monomer showed a similar phenotype in cells, but *in vitro* showed a difference in ability to bind to PI(3)P, suggesting that some of the negative regulation of DNAJC13_FL_ that occurs in cells is lost in the detergent lysate. While future studies will be required to determine if DNAJC13 oligomerizes in cells, recent work on RME-8 identified a series of self-interactions which could allow for oligomerization (Norris *et al*., 2022). These interactions between RME-8 domains were first mapped by pulldown and yeast two-hybrid screens as occurring between the J domain and a C-terminal region of RME-8 (1650-2279), and the residues in the C-terminus were later mapped to D1657 and E1962 in repeating motifs called IWNs (Shi *et al*., 2009; Norris *et al*., 2017). While these C-terminal control points in RME- 8 are different from those we identify in human DNAJC13, it points toward a general model of the DNAJC13/RME-8 C-terminus performing a regulatory role.

### DNAJC13 C-terminal Tail as a Disordered Regulatory Region

Intrinsically disordered regions (IDRs) often play regulatory roles in protein function (Fenton *et al*., 2023). Here we use two predictors of structural disorder, AF and JRonn, to demonstrate that the C-terminal tail of DNAJC13 is likely to be disordered. We then identified a novel and conserved motif we refer to as YLT1, consisting of Y2205, L2206, T2207, as a key negative regulator of DNAJC13 localization to endosomes in cells and ability to bind to PI(3)P *in vitro*. Interestingly, we identify a second occurrence of the YLT sequence (YLT2; Y2215, L2216, T2217), which upon mutation also results in enhanced localization of DNAJC13 to endosomes, albeit much weaker than mutation of the YLT1 sequence. Another feature of IDRs is that they are often the target of post-translational modification, and the C-terminal tail of DNAJC13 is in fact overrepresented in residues able to be phosphorylated (13 residues, 29% of residues) (Fenton *et al*., 2023). While future studies will be necessary to determine what the YLT1 motif interacts with, it appealing to consider a model in which the DNAJC13 C-terminal tail makes autoinhibitory contacts within DNAJC13 itself, and that this interaction can be further regulated by dynamic phosphorylation and/or protein binding.

Notably, endogenous DNAJC13 and RME-8 are primarily localized to endosomes and this phenotype is distinct from the largely cytoplasmic localization of the overexpressed GFP- DNAJC13_FL_ that we see here (Zhang *et al*., 2001; Fujibayashi *et al*., 2008; Freeman *et al*., 2014; Novy *et al*., 2024). The relative distribution of GFP-DNAJC13_FL_ between endosomes and the cytoplasm may have a cell-type dependent component as some reports show GFP-DNAJC13_FL_ as primarily cytoplasmic while others show it more localized to vesicles (Fujibayashi *et al*., 2008; Freeman *et al*., 2014; Xhabija and Vacratsis, 2015). Importantly, one study that identified GFP- DNAJC13_FL_ as primarily localized to endosomes used a pre-fixation digitonin treatment, which specifically reduces cytoplasmic signal (Liu *et al*., 2001; Fujibayashi *et al*., 2008). In our hands the largely cytoplasmic phenotype GFP-DNAJC13_FL_ was useful as it allowed us to perform a structure/function analysis of domains in DNAJC13 which negatively regulate its localization to endosomes. However, it is likely that our study did not capture all mechanisms controlling DNAJC13 localization to endosomes including interactions with other binding partners (e.g. SNX1, FAM21) which–while not necessary for its endosomal localization–may help stabilize DNAJC13 on endosomes (Harbour *et al*., 2012; Jia *et al*., 2012; Helfer *et al*., 2013; Freeman *et al*., 2014; Xhabija and Vacratsis, 2015; Dostál *et al*., 2023).

### DNAJC13 proteomics: HSC70 and co-chaperones

In our quantitative proteomics examining DNAJC13_FL_ interactors, we found both HSC70, a member of the Hsp70 family, and also known HSC70 co-chaperones. While HSC70 is a known interactor of DNAJC13, it had not been previously noted that DNAJC13 would co-purify with HSC70 co-chaperones (Chang *et al*., 2004; Girard *et al*., 2005). Broadly, the Hsp70 family of proteins are ATPases responsible for unfolding/refolding of proteins and disassembly of protein complexes and require several co-chaperones which function at different stages of its catalytic cycle. First, the J domain containing co-chaperone brings a client (e.g. substrate) to the HSC70 substrate binding domain and the HPD containing J domain binds the HSC70 nucleotide binding domain, stimulating HSC70 ATPase activity (Bracher and Verghese, 2015). This induces the conformational change responsible for client unfolding or complex disassembly (Bracher and Verghese, 2015). Then, a nucleotide exchange factor (NEF) co-chaperone binds HSC70 and stimulates the exchange of ADP for ATP, which primes HSC70 for another round of activity. Lastly, other HSC70 co-chaperones exist to slow down its catalytic cycle or target clients for degradation (Bracher and Verghese, 2015).

In our interaction proteomics, we found three members of the Hsp70 family; HSPA8/HSC70, the known interactor of DNAJC13 (Chang *et al*., 2004; Girard *et al*., 2005; Ryu *et al*., 2020); HSPA1A/HSP70, the heat shock inducible paralogue (Bilog *et al*., 2019); and HSPA9, a mitochondrial paralogue (Luo *et al*., 2010). Capture of the mitochondrial paralogue is likely a result of detergent-based purification of DNAJC13, as it would not normally be at the right place for interaction with DNAJC13. Interestingly, we also identified two NEFs from different families: HSPA4 (Hsp110 family) (Kaneko *et al*., 1997), and BAG2 (BAG family) (Arndt *et al*., 2005). As J domains and NEFs both bind the Hsp70 nucleotide binding domain and promote opposite ends of its ATPase cycle, it is curious why we would capture NEFs in our DNAJC13 proteomics. One possible explanation was suggested by recent unbiased proteomics which sought to globally characterize HSP70 and HSC70 co-chaperones and client proteins. In this study, they identified endogenous DNAJC13 as a unique type of J domain containing protein because, in addition to being a specific co-chaperone of HSC70, it was also found as a potential HSC70 client protein (Ryu *et al*., 2020). Thus, it is possible that the presence of NEFs in our proteomics support a model in which DNAJC13 is both a co-chaperone and client of HSC70.

Lastly, we also identified STUB1 (also known as CHIP) in our DNAJC13 proteomics.

STUB1 is a co-chaperone that binds Hsp70 C-terminal domain and has dual functions of slowing down HSC70 ATPase activity as well as being an E3-ligase that can ubiquitinate clients that have failed refolding (Meacham *et al*., 2001; Stankiewicz *et al*., 2010). While our proteomics cannot distinguish which co-chaperones are binding the same HSC70 protein, it is interesting to note that BAG2 and CHIP can exist in a multi-member complex with Hsp70s, where BAG2 inhibits CHIP binding to other members of the ubiquitin ligase machinery, and thus BAG2 inhibits ubiquitin-dependent client degradation (Arndt *et al*., 2005; Dai *et al*., 2005). Together, we identify that DNAJC13 interacts not just with HSC70 but active HSC70 complexes including those bound to several types of co-chaperones. Our findings support the previously identified interaction between HSC70 and DNAJC13/RME-8 and suggest that, in addition to functioning as a co-chaperone, DNAJC13 may also be a client of HSC70 (Chang *et al*., 2004; Girard *et al*., 2005; Ryu *et al*., 2020).

### Targets of the DNAJC13 J domain

While J domain-containing proteins are often thought of in terms of proteostasis, the role of J domains in membrane trafficking has been best studied in endocytosis where auxilin is involved in uncoating clathrin coated vesicles (Eisenberg and Greene, 2007). In this mechanism, auxilin binds clathrin, recruits HSC70, and stimulates the ATPase activity of HSC70 through the catalytic triad HPD in its J domain (Morgan *et al*., 2001; Eisenberg and Greene, 2007). The current model of DNAJC13/RME-8 function is that it recruits HSC70 to disassemble proteins on the endosomes including specific targets like clathrin and SNX1 (Girard *et al*., 2005; Popoff *et al*., 2009, 2009; Xhabija and Vacratsis, 2015). While it is worth noting that these experiments used loss of overall DNAJC13 as a proxy for J domain activity, similar effects on endosomal protein function were observed upon manipulation of HSC70 function (Zhang *et al*., 2001; Chang *et al*., 2004; Popoff *et al*., 2009; Shi *et al*., 2009).

Our findings add to this model and suggest an additional target of the DNAJC13/HSC70 interaction: DNAJC13 itself. Specifically, our unbiased proteomics shows that HSC70 is the primary interactor of overexpressed DNAJC13_FL_ and that loss of DNAJC13’s ability to bind and stimulate the ATPase activity of HSC70 (DNAJC13hpd) results in increased DNAJC13 endosomal localization in cells and PI(3)P binding *in vitro*. Additionally, we identify NEFs—which promote the opposite part of the HSC70 catalytic cycle as J domains—in our proteomics and recent unbiased proteomics identified endogenous DNAJC13 as a co-chaperone and potential client of HSC70. Together these findings support a model that DNAJC13 is an “atypical” J domain containing protein and may be a target of its own J domain activity. Combined with our findings about an oligomerization requirement for DNAJC13_PH-like_ domain to associate with PI(3)P/endosomes, and the recent proposal that *C. elegans* RME-8 oligomerizes, it is intriguing to speculate that HSC70 regulates DNAJC13 localization and function through disassembly of DNAJC13 oligomers (Norris *et al*., 2022).

### DNAJC13, WASH complex, Clathrin, and Endosomal Clustering

We observed that mutation of the catalytic triad of the DNAJC13 J domain, particularly when combined with truncation of the DNAJC13 C-terminal tail, resulted in a higher propensity for DNAJC13-positive endosomes to cluster together and collapse into a perinuclear region. This observation is strikingly similar to what was observed upon loss of function of two DNAJC13 interacting proteins/complexes: the WASH complex and clathrin heavy chain. Specifically, loss of WASH complex function (knockout of the WASH1 subunit) or disruption of clathrin function (overexpression of the dominant negative hub domain of clathrin heavy chain) results in a redistribution of EEA1 positive endosomes from distributed throughout the cell to tightly clustered and collapsed perinuclear region (Bennett *et al*., 2001; Gomez *et al*., 2012). The mechanism by which disruption of WASH, clathrin, or DNAJC13 causes endosomal collapse is unknown, although DNAJC13 has been shown to regulate both clathrin and WASH, thus functionally linking these three proteins/complexes (Chang *et al*., 2004; Shi *et al*., 2009; Freeman *et al*., 2014; Xhabija and Vacratsis, 2015; Novy *et al*., 2024). It is interesting to note that recent work has linked the WASH complex to the dynein/microtubule system, which promotes endosomal translocation to the perinuclear region (Fokin and Gautreau, 2021; Fokin *et al*., 2021). Thus, our data demonstrate that similar to clathrin and WASH, disruption of endogenous DNAJC13 results in disruption of endosome distribution in cells.

Together, our study examined how human DNAJC13, a protein important in endosomal sorting, is regulated. We identify that DNAJC13 localization to endosomes is controlled by the low affinity of its PH-like domain for PI(3)P, which can be overcome by oligomerization, and the negative regulation promoted by its J domain and C-terminal tail. Future studies will be important in showing how these novel control points integrate cellular signals to tune DNAJC13 function on endosomes and thereby control efficient cargo sorting into the recycling and degradative pathways.

## Acknowledgements

We thank the rest of the Lobingier Lab (T. Weishaar and A. Dagunts) for advice and feedback on this paper. We thank Kiyotoshi Sekiguchi for providing GFP-DNAJC13. We thank the OHSU Proteomics Shared Resource core for assistance with TMT labeling, mass spectrometry and proteomics data analysis (A. Reddy and P. Wilmarth; supported by the National Institutes of Health under core grants P30EY010572, P30CA069533, and S10OD012246). This work was carried out with the help of other core facility resources: OHSU Flow Cytometry Core (P. Canaday), the OHSU Advanced Light Microscopy Core (RRID:SCR_009961, F. Kelly and S. Kaech Petrie). B.T.L was supported by GM137835 and OHSU startup funds. H.A. was supported by T32GM142619.

## Materials and Methods

### Chemicals

From Corning, DPBS without Calcium or Magnesium (Corning, 21-031-CV) and DPBS with Calcium and Magnesium (Corning, 21-030-CM). From Sigma-Aldrich, Bovine Serum Albumin (A7030) was dissolved in DPBS with Calcium and Magnesium and filtered before use. For cell fixation for microscopy, 16% paraformaldehyde ampules were purchased from Invitrogen (Thermo Scientific, 28906) and diluted to 4% in DPBS with Calcium and Magnesium immediately before use.

### Antibodies

From Cell Signaling, mouse anti-EEA1 (Cell Signaling, 48453S) and mouse anti-GFP (55494S), rabbit anti-GM130 (Cell Signaling, 12480T, for imaging). From Novus Biologicals, rabbit anti-GFP (Novus Biologicals, NB600-308, for imaging). From Takara Biosciences, mouse anti-GFP (Clontech Labs 3P 632381, for western blot). Secondary antibodies for imaging from Invitrogen – goat anti-mouse AF488 (Thermo Scientific, A11029), goat anti-rabbit AF488 (Thermo Scientific, A32731), Goat anti-mouse AF647 (Thermo Scientific, A21235), goat anti-rabbit AF647 (A32733). Secondary antibodies for western blotting from Bio Rad goat anti-mouse StarBrite 700 (Bio-Rad, 12004158).

### Structural prediction

The AlphaFold2 structural prediction was downloaded from the AlphaFold Protein Structural Database (https://alphafold.ebi.ac.uk/entry/O75165) (Varadi *et al*., 2022) and visualized in Pymol. For AlphaFold3 structural prediction, the sequence for human DNAJC13 (Uniprot O75165) was input into the DeepMind AlphaFold3 server (https://golgi.sandbox.google.com/) with a random seed (Abramson *et al*., 2024). All models were downloaded and viewed separately in Pymol, where the final 73 residues were each given a score of 1 for unstructured and 0 for structured. The average of the 5 models is shown in Figure S1A.

For JRonn disorder prediction, the sequence for DNAJC13 (O75165) was opened in Jalview (Waterhouse *et al*., 2009) and the C-terminal 257 amino acids were run through the homology-based secondary structure JPred algorithms, including the JRonn disorder predictor algorithm.

### Sequence conservation

To assess the C-terminus for sequence conservation, all vertebrate (plus *D. melanogaster* and *C. elegans*) orthologues for human DNAJC13 were downloaded from the Ensembl database (Harrison *et al*., 2024) as a multiple sequence alignment. This alignment was opened in Jalview, trimmed to show only sequences aligning with the human C-terminal tail and relative conservation score was calculated(Waterhouse *et al*., 2009).

### DNA constructs

All plasmids were verified either via Sanger sequencing of several reads or whole plasmid nanopore sequencing. pEGFP-DNAJC13 was a gift from the Sekiguchi group(Fujibayashi *et al*., 2008). Upon sequencing of our construct, we noticed a nonnative sequence on the C-terminus (HRPLPGSTGSR) and removed this sequence by re-cloning the native sequence into the parental pEGFP-C1 vector between restriction sites KpnI and BamHI and the resulting construct is what we refer to as DNAJC13_FL_. To create the C-terminally tagged DNAJC13_FL_, GFP was PCR’d and inserted to the C-terminus of pEGFP-DNAJC13_FL_ using NEBuilder (New England Biologicals, E2621L) to insert at the BamHI site. After successful insertion, the N-terminal GFP was removed by digestion with AgeI and KpnI, and NEBuilder to stitch the plasmid back together with a new start codon, creating pEGFP-DNAJC13_FL_-ctGFP. This construct begins with the linker between the original N-terminal GFP and DNAJC13 (GGGSGGGS).

PCR, digestion and ligation with KpnI and BamHI were again used to copy specific regions and re-insert into the parental pEGFP-C1 vector for truncated protein DNAJC13_2198t_ from DNAJC13_FL_. To perform the alanine scanning of the c-terminal tail, double stranded gBlocks from IDT were obtained containing the mutant sequences as well as homology arms for assembly with NEBuilder after digestion of pEGFP-DNAJC13_2198t_. To mutate the DnaJ domain residues (HPD) to alanine, a shorter construct encoding residues 1-1927 of DNAJC13 was cloned into pEGFP-C1 vector between KpnI and BamHI. Next, a gBlock from IDT was obtained encoding for a fragment of DNAJC13 with the HPD residues mutated to alanine and inserted between internal cut sites BlpI and PshAI with NEBuilder. Next, the C-terminus encoding 1927- 2198 or 1927-end was copied via PCR and inserted into the end of the truncated, hpd mutant construct after the BamHI site using NEBuilder, creating DNAJC13hpd and DNAJC13_2198t(hpd)_.

Truncated proteins DNAJC13_t347_, DNAJC13_t347(ylt1)_, DNAJC13_t347(hpd)_, and DNAJC13_351t_ were created by PCR of the region from DNAJC13_FL_, or DNAJC13_ylt1_ or DNAJC13hpd for the respective mutants, and reinsertion (via NEBuilder for DNAJC13_t347_ constructs, and classical linear ligation for DNAJC13_351t_) into the parental pEGFP-C1 vector between KpnI and BamHI. To add dimerizing and tetramerizing domains to 351t, dimerizing and tetramerizing motifs were codon corrected from the original sequence for bacterial expression (Khairil Anuar *et al*., 2019) for human cell expression and ordered as gBlocks from IDT with homology overlaps for cloning into pEGFP-DNAJC13_351t_ at the BamHI site.

### Cell culture

FLP-In-293 (Thermo Scientific, R75007) cells were purchased from Thermo Fisher Scientific and HeLa (ATCC, CCL-2) were purchased from ATCC. Both were grown in DMEM (Thermo Fisher Scientific, 11965-092) supplemented with 10% FBS, at 37°C and 5% CO2.

### Plasmid transfection

For microscopy, flow cytometry, and western blot experiments, HeLa cells were plated at 50% confluence in dishes for the respective experiment. The next day they were transfected using Lipofectamine-2000 (Thermo Scientific, 11668019) and OptiMEM (Gibco, 31985088). DNA, lipofectamine, and OptiMEM was scaled for the experiment and DNA/lipofectamine-200 used depended on the length of the construct, with bigger constructs having more DNA/lipofectamine and smaller constructs less. DNAJC13_FL_, DNAJC13hpd, and triplet scanning mutants were all transfected at 1.25x amounts, while DNAJC13_2198t_, DNAJC13_2198t(hpd)_, DNAJC13_t347_, DNAJC13_t347(ylt1)_, and DNAJC13_t347(hpd)_ were transfected at 1x amounts, and DNAJC13_351t_, DNAJC13_351t_-dimer and DNAJC13_351t_-tetramer were transfected at .75x amounts.

Imaging experiments using 8 well imaging dishes (Thermo Scientific, 155409) were transfected with Lipofectamine-2000 (0.643 µL 1x) and DNA (300 ng 1x) in OptiMEM (50 µL). Flow cytometry experiments were performed in 12 well dishes and were transfected with Lipofectamine-2000 (1.875 µL 1x) and DNA (875 ng 1x) in OptiMEM (400 µL). Western blot expression experiments were performed in 6 well dishes and were transfected with Lipofectamine-2000 (5.14 µL 1x) and DNA (2400 ng 1x) in OptiMEM (400 µL). Fixed microscopy experiments were performed in 24 well dishes containing #1.5 thickness round cover slips (Harvard Apparatus, 64-0712) coated in 1:100 Poly-L-Lysine (Sigma-Aldrich, P8920-100ML) and were transfected with Lipofectamine-2000 (1.22 µL 1x) and DNA (570 ng 1x) in OptiMEM (120 µL).

For PIP binding studies and CoIP proteomics studies, FLP-In-293 cells were used instead of HeLa cells. For PIP binding studies, they were plated at 40% confluence in T25s. The next day they were transfected with Lipofectamine-2000 (27.3 µL) and DNA (13.3 ng) in OptiMEM (1 mL). For CoIP proteomic studies, they were plated at 20% confluence in T182s. The next day they were transfected with 81.5 µL Lipofectamine-2000 (81.5 µL) and DNA (81.5 ng) in OptiMEM (1 mL).

### Flow cytometry for expression

One day after transfection with GFP-DNAJC13 constructs, cells were washed with DPBS without Ca/Mg and lifted in TrypLE (Gibco, 12604021) and resuspended in Flow Buffer (DPBS+Ca/Mg + 1% BSA). Cells were analyzed using a Beckman Coulter CytoflexS. For each experiment, 10,000 counts were taken after discrimination of cells (forward vs side scatter) and singlets (forward scatter vs forward scatter width). Data was then reanalyzed via FlowJo to gate for cells and singlets and assess the geometric mean of the FITC-A channel (488 nm laser, 525/40 nm filter).

### Live cell microscopy

One day after transfection with GFP-DNAJC13 constructs, cells were treated with 1:4000 Invitrogen CellMask Deep Red Plasma membrane stain (Thermo Scientific, C10046) and 1:500 Pierce Hoechst-33342 DNA stain (Thermo Scientific, 62249) diluted in pre-equilibrated Fluorobrite (Thermo Scientific, A1896701). After 10 minutes in the incubator, media was replaced with fresh, pre-equilibrated Fluorobrite and moved to the imaging incubator (35°C) on a Nikon spinning disk confocal microscope (Yokogawa CSU-W1 on a Nikon TiE). Cells were imaged under a 100x oil immersion objective (1.49 NA, Apochromat TIRF, 12 mm working distance) with the blue channel (405 nm laser, 445/50 nm filter), green channel (488 nm laser, 525/36 nm filter), and far-red channel (640 nm, 700/75 nm filter). Each construct was imaged over three biological replicates, taking 6-12 images per construct each replicate.

### Blinded analysis of phenotype

All images had cells manually sectioned and ROIs were saved in FIJI-ImageJ. Images and ROI sets for all constructs to be blinded (GFP-DNAJC13_FL_, GFP-DNAJC13_2198t_, GFP-DNAJC13_ylt1_, GFP-DNAJC13_hpd_, and GFP-DNAJC13_2198t(hpd)_) were renamed to randomized numbers.

Individual cells were scored into two initial phenotypes as follows: cytoplasmic if they had a bright cytoplasmic background, containing some localized puncta; and localized if they had a dim cytoplasmic background and bright punctal localization. The localized phenotype was further dissected into two: distributed if the endosomes were spread across the cell; and clustered if endosomes were largely confined to one or two contiguous structures.

### GFP signal accumulation metric

Cells were manually sectioned and analyzed for maximal and median pixel intensity of the green channel in FIJI-ImageJ. For samples that had blinded phenotypic analysis performed, ROIs were the same ones used in both analyses to allow for direct comparison of phenotype and quantitative metrics. GFP signal accumulation was found by dividing the maximal pixel intensity by the median pixel intensity. All healthy cells imaged over the three biological replicates were included as individual points for analysis, and the mean scores from each replicate were compared in statistical analysis as a SuperPlot.

### Fixed microscopy

One day after transfection, coverslips were washed with DPBS+Ca/Mg before fixing for 20 minutes with 4% paraformaldehyde while rocking at RT. Cells were rinsed 3x with DPBS+Ca/Mg, blocked and permeabilized for 30 minutes, rocking at RT with Imaging Block Buffer (DPBS+Ca/Mg+4% BSA+0.1%TritonX), then incubated with primary antibodies overnight, rocking at 4°C (1:1000 rabbit anti-GFP and 1:500 mouse anti-EEA1, or 1:1000 mouse anti-GFP (Cell Signaling) and 1:1000 rabbit anti-GM130, diluted in Imaging Block Buffer). The next day, cover slips were rinsed 3x with DPBS+Ca/Mg, incubated with secondary antibodies (1:2000 anti-Mouse-488 & anti-Rabbit-647 or 1:2000 anti-Rabbit-488 & anti-Mouse-647 in Imaging Block Buffer) for 1 hour rocking at RT before being washed 3x with DPBS+Ca/Mg and mounted on fresh glass slides with ProLong Diamond + DAPI (Thermo Scientific, P36962).

At least one day after mounting, cells were imaged using the same Nikon spinning disk confocal microscope as used for live microscopy. On three separate biological replicates for all constructs analyzed with fixed microscopy, 5 fields of view were imaged with Z-stacks covering whole cells, a representative example of a single z-plane is shown.

### SDS-PAGE sample preparation for construct expression

For analyzing expression of GFP-tagged constructs, one day after transfection, cells were washed once with DPBS and lifted with TrypLE. Cell pellets were collected and lysed on ice for 10 minutes with 250 µL RIPA Buffer (50 mM Tris pH 7.4, 150 mM NaCl, 1% TritonX, 0.5% sodium deoxycholate, 0.1% sodium dodecyl sulfate) with HALT protease inhibitor cocktail (Thermo Scientific, 78430). Cells were further lysed via sonication (1s on/3 s off, 3 cycles at 35% amplitude). Lysates were then clarified at 10,000 x g for 10 minutes at 5°C and a sample was combined with 4x SDS PAGE Sample Buffer (250 mM Tris, pH 6.8, 40% glycerol, 8% SDS, bromophenol blue) + beta-mercaptoethanol and heated at 95°C for 5 minutes.

### Western blotting protocol

Samples were loaded along with ladder (Bio-Rad; 1610363, 1610373, 1610377; or GoldBio, P007) onto gradient Bio-Rad 4-20% polyacrylamide SDS-PAGE gels containing StainFree total protein stain (Bio-Rad, 456-8095) and run at 125V in SDS-PAGE running buffer (250.1 mM Tris, 1.924 M glycine, 0.0347 M SDS) until dye front ran off the gel. StainFree total protein stain was activated on a Bio-Rad ChemIDoc Imaging System and imaged before transfer onto nitrocellulose with the Bio-Rad TurboBlot Transfer system (Bio-Rad, 1704150). Blots were then blocked in Bio-Rad EveryBlot Blocking Buffer (Bio-Rad, 12010020) for ∼90 min rocking at RT, then primary antibody (Takara Biosciences mouse-anti-GFP, 1:1000) was diluted in Western Blot Antibody Buffer (1xTBS pH 7.4 + 5% BSA + 0.1% TritonX) and rocked at 4°C overnight.

Blots were washed four times with PBST (DPBS+0.1%TritonX). Bio-Rad StarBrite secondary antibody (1:3000, diluted in PBST) were incubated for 1 hour rocking at RT before being washed four times with PBST and imaged on the Bio-Rad ChemIDoc.

### Coimmunoprecipitation proteomics sample preparation and processing

One day after transfection with GFP or GFP-DNAJC13, HEK293 cells were lifted with TrypLE and quenched with DMEM. A small sample was resuspended in Flow Buffer and taken to flow cytometry (see *Flow Cytometry for expression*). The geometric means of FITC fluorescence after gating for cells and singlets were used calculate normalization factors. Cells were then lysed in 3.6 mL CoIP Lysis Buffer (10 mM Tris pH 7.5, 150 mM NaCl, 1% TritonX) with HALT protease inhibitor cocktail. After 10 minutes on ice, lysates were diluted with CoIP Lysis Buffer to normalize GFP loading onto resins. 20 µL of Chromotek GFP-Trap resin slurry (Chromotek, gta-20) were equilibrated with CoIP Lysis Buffer and 1.2 mL of normalized lysates were loaded onto each resin and bound for 1 hour at 4°C while rocking. Resins were then washed 3x with CoIP Lysis Buffer and eluted twice by boiling with 192 µL 5% SDS. Eluates were combined and frozen before processing for proteomics.

Eluates were thawed, buffer was added (TEAB to 50 mM from 1 M stock), reduced with 22 mM DTT, cysteines methylated with 40 mM iodoacetamide (Thermo Scientific, A39271). Protein was then purified and proteolyzed on-column with Trypsin/LysC (Thermo Scientific, A40007) on Protifi S-trap micro columns (Protifi, C02-micro-10) according to manufacturer’s protocol. In brief, eluates were acidified to pH of ∼1 with Phosphoric acid, diluted in 6 volumes of S-Trap Protein Binding Buffer (90% aq methanol, 100 mM TEAB, pH 7.5) before loading on S-Trap columns. Columns were extensively washed with S-Trap Protein Binding Buffer before overnight digestion at 37°C with 2 µg Trypsin/LysC mix. The next day, columns were rehydrated with 20 µL 50 mM TEAB (pH 7.5) and digested peptides were eluted in three separate eluates consisting of; 1) 40 µL 50 mM TEAB; 2) 40 µL 0.2% formic acid and; 3) 40 µL 50% ACN, 0.2% formic acid. Eluates were combined and lyophilized. Peptides were resuspended in 100 µL 30% ACN, a sample was taken for quantification with the Pierce Peptide Assay (Thermo Scientific, 23275), and the rest was lyophilized. Peptides were resuspended, and a normalized amount (4.4 µg) was taken for labeling with TMTpro 10-plex Label Reagents (Thermo Scientific, A52047), quenched and dried. 4 µg labeled peptide was pooled, quenched with 0.5% final hydroxylamine, dried down and resuspended in 40 uL 10 mM ammonium formate (pH 10).

These were analyzed by LC-MS on a Dionex Ultimate HPLC operating in 2D mode (mobile phase: 20-90% ACN pH 9, flow rate: 3 uL /min; 7.5-30% low pH; flow rate: 300 nL/min) coupled to the Orbitrap Fusion Tribrid mass spectrometer using the SPS MS^3^ scan mode for TMT quantification (data-dependent MS2 scans using dynamic exclusion, resolution: 120K).

For analysis, the PAW pipeline (Wilmarth *et al*., 2009) using Comet search engine (version 2016.03) (Eng *et al*., 2013) were used to extract spectra, search against a Uniprot human database with added contaminants and eGFP (downloaded October, 2020, 20605 protein entries plus; eGFP, 174 common contaminant sequences, and sequence-reversed decoys). Comet was configured with static cysteine alkylation (+57.0215 Da), static TMTpro reagent modifications (+304.2071 Da) on lysines and peptide N-termini, variable oxidation of methionine, a parent ion mass tolerance of 1.25 Da, a fragment ion mass tolerance of 1.005 Da and full tryptic digest with a maximum of two missed cleavages. Identified peptides were then filtered using a reversed-sequence decoy strategy (Elias and Gygi, 2007) to control peptide spectrum match false discovery at an FDR of 1%. At least two unique peptides were required for positive identification of a protein from the data. A list of inferred proteins and TMT reporter ion intensities per channel was exported for statistical analysis, where intensities were compared between groups using the Bioconductor package edgeR (Robinson *et al*., 2010) after trimmed mean of *M*-value normalization (Robinson and Oshlack, 2010) in RStudio. Multiple-testing corrections and calculation of FDRs was performed within edgeR using Benjamini-Hochberg method, and hits were selected based on an FDR of <1%.

### Phosphatidylinositol phosphate (PIP) binding studies

Protocols adapted from (Xhabija and Vacratsis, 2015), in brief; HEK293 cells were seeded in a T25 at 40% confluence, 24 hours later, they were transfected with GFP-DNAJC13 constructs. The next day, cells were lifted with TrypLE, quenched with DMEM, a small sample was resuspended in Flow Buffer and analyzed on a Beckman Coulter Cytoflex S (see *Flow Cytometry for expression*). Using FITC-A geometric mean to normalize GFP loading, cells were lysed in a varying amount of PIP Lysis Buffer (50 mM Tris, pH 7.4, 76 mM NaCl, 1% TritonX, 10% glycerol, 2 mM EGTA) with HALT protease inhibitor cocktail, on ice by sonication (1s on/3 s off, 7 cycles @35% amplitude). A portion of lysate was then clarified by centrifugation (15,000 x g, 10 min, 4°C). A sample of clarified lysate was taken for western blot analysis and 250 µL loaded onto phosphoinositide decorated resins (50 µL slurry) - PIP (Echelon Biosciences, P- B001) and PI(3)P (Echelon Biosciences, P-B003A), pre-equilibrated in PIP Lysis Buffer. Lysates were bound for 2 hours on a rotisserie at 4°C. Resins were then washed three times in PIP Wash Buffer (10 mM HEPES pH 7.4, 150 mM NaCl, 0.25% TritonX) before elution with 2xSDS PAGE Sample Buffer (diluted from 4x in PIP Wash Buffer) at 70°C for 10 minutes.

### Statistical analysis and reproducibility

Statistical analysis was performed in Prism (GraphPad) or published software for proteomics (PAW_Pipelinev0616a7f). All experiments except the proteomics come from at least three biological replicates, which comes from two. Additionally, DNAJC13_t347_ PI(3)P binding was only performed twice while the other samples in the set were performed in triplicate (Figure 4B).

Plotted data are represented as individual biological replicates, or as SuperPlots with the means of at least three biological replicates, as well as data from individual cells across replicates, where replicate averages were compared for statistical analysis (Lord *et al*., 2020). Expression western blots were performed on three separate experiments for all constructs and a representative example is shown. All measurements were taken from distinct samples, except as follows: DNAJC13_FL_ GFP signal accumulation data is used as control for comparison in 1F, 2C, 3D, 4F; DNAJC13_FL_ Flow cytometry data is reused between 1B, S2B, S3B; and DNAJC13_FL_ flow cytometry data is reused between S4C, S5C. Statistical test performed is noted in each figure legend, unpaired two-tailed t-test (1F), unpaired one-way ANOVA with Dunnett’s multiple comparison’s corrections (2C, 3D, 4B, 4D, 4F), paired one-way ANOVA with Tukey’s multiple comparison’s corrections (5C), or unpaired one-way ANOVA with Tukey’s multiple comparison’s corrections (5E). P values are represented as: ns if P>0.05, * if P<= 0.05, ** if P <= 0.01, *** if P <= 0.001, and **** if P <= 0.0001.

### Software and code

Data were collected with the following software: flow cytometry (Beckman CytExpert, v2.4), western blot (Bio-Rad Image Lab Touch v2.4.0.03 and FIJI-ImageJ v2.14.0/1.54f), and microscopy (Nikon Elements v4.51.01 (Build 1146)). Data were analyzed with the following software: statistical analysis and graphing (GraphPad Prism v10.3.1), flow cytometry (FlowJo v10.10.0), proteomics (PAW-Pipeline v0616a7f https://github.com/pwilmart/PAW_pipeline with Comet search engine v2016.03, statistical analysis in RStudio v2023.09.01 build 494 with edgeR v4.0.16) , and microscopy (FIJI-ImageJ v2.14.0/1.54f). JRonn modeling and conservation analysis were performed in Jalview (v 2.11.4.1). Structural analysis of models was performed in Pymol (Schrodinger Pymol v 2.5.7).

## Data availability

All data generated and analyzed in this study are included as figures or supplementary information. The human proteome was downloaded from the Uniprot human protein database at https://www.uniprot.org/proteomes/UP000005640. Raw and analyzed proteomics data have been deposited at the ProteomeXchange Consortium via the PRIDE partner repository with the dataset identifier PXD058964. Source data are provided within this paper.

## Supporting information

This article contains supporting information.

## Author contributions

B.T.L. directed the study. H.A. performed primary experimentation and preparation of figures.

H.A. performed proteomics coimmunoprecipitation workup and analysis. Constructs were cloned by all authors. B.N. performed some initial microscopic observations. E.H. optimized PI(3)P binding experiments. H.A. wrote the manuscript with support from B.T.L. and critical feedback from other authors.

## Funding and additional information

This work is supported by the National Institute of Health (GM137835 to B.T.L, T32GM142619 to H.A.). B.T.L. is also supported by OHSU startup funds.

---

**Figure S1.**
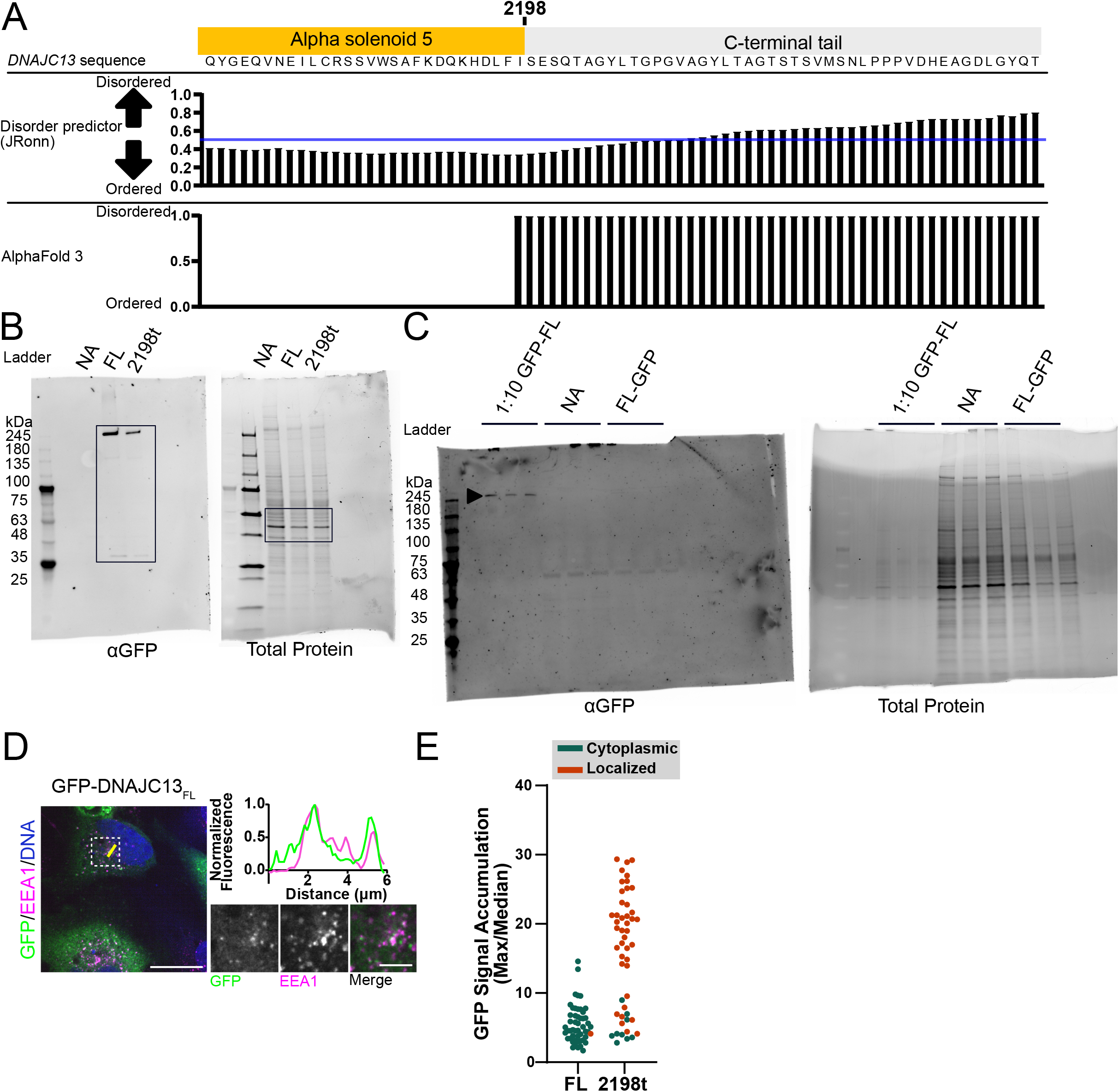
***A***, Structural prediction for the C-terminus of DNAJC13 (sequence, above), with JRonn disorder prediction (middle) and summary from five AlphaFold3 structural predictions (bottom). ***B***, Uncropped anti-GFP western blot (left) and total protein stain gel (right) from Figure 1B; cropped area shown in the black box. ***C***, Western blot (anti-GFP, left) and total protein stain (right) of three replicates of HeLa cells transfected with DNAJC13_FL_ (at a 1:10 dilution of a standard load) or DNAJC13_FL_-ctGFP (undiluted), and a nontransfected control. ***D***, Fixed immunofluorescent microscopy image of GFP-DNAJC13_FL_ expressed in HeLa cells. Imaged with anti-GFP (green), endosomal marker EEA1 (magenta), and DAPI DNA stain (blue) with insets shown to the right (scale bar = 20 µm, 5 µm in inset), (representative example from n=3 biological replicates). A line-scan (yellow line) showing normalized fluorescent intensity of GFP (green) and EEA1 (magenta) signal are plotted along the line (right). ***E***, Signal accumulation metric data (from Figure 1F), correlated by color to the blinded phenotypic scoring (from Figure 1G).

**Figure S2.**
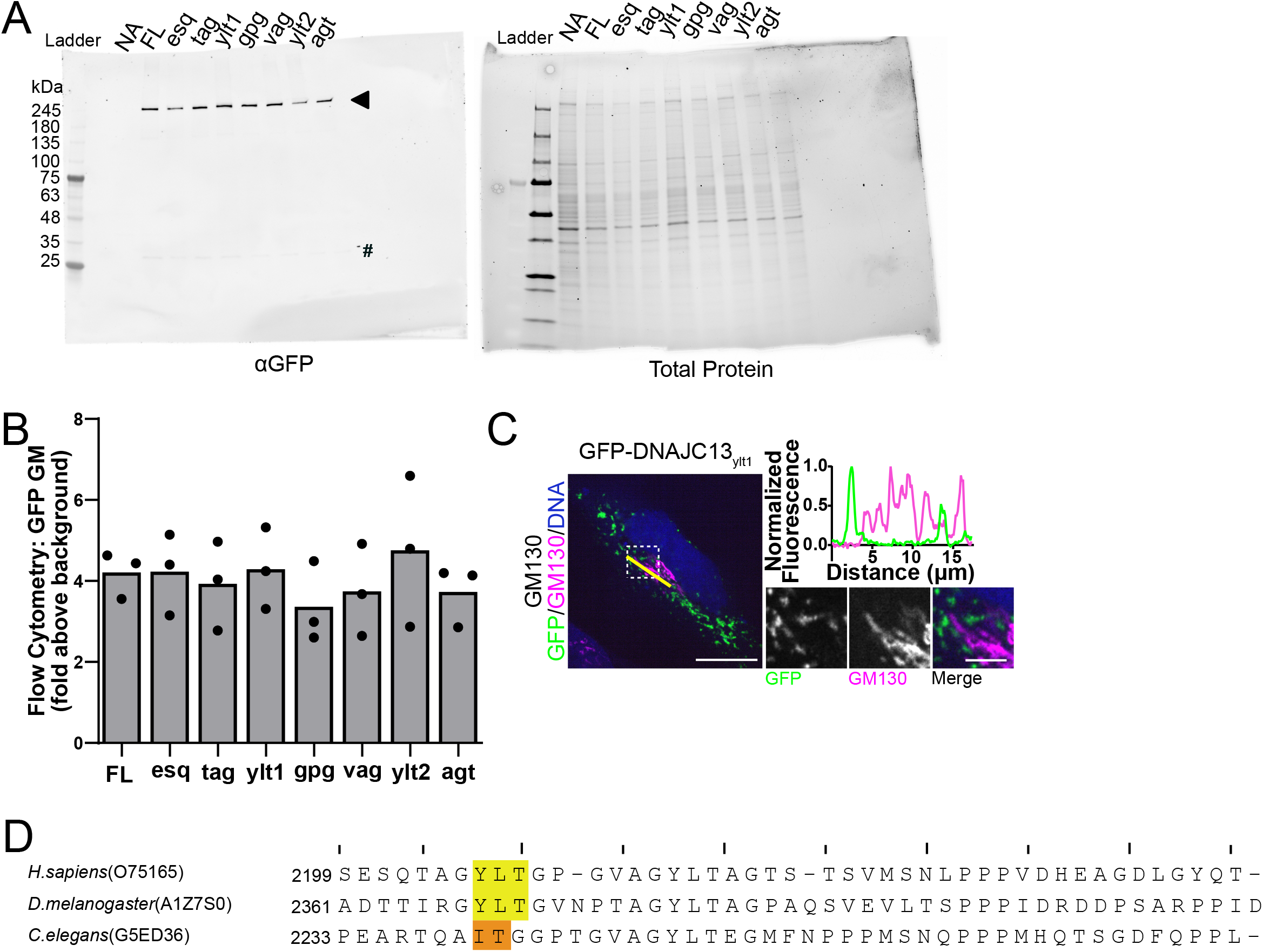
***A***, Representative western blot (anti-GFP, left) and total protein stain gel (right) of HeLa cells transfected with DNAJC13_FL_ or triplet scanning mutants, and a nontransfected control (n=3 biological replicates). The arrowhead marks GFP-DNAJC13 and the # marks free GFP. ***B***, Flow cytometry-based expression analysis of constructs expressed in HeLa cells, assessed by geometric mean of GFP channel, displayed as fold above background signal from untransfected cells (n=3 biological replicates). ***C***, Fixed immunofluorescent microscopy image of GFP-DNAJC13_ylt1_ expressed in HeLa cells. Imaged with anti-GFP (Green), Golgi marker anti- GM130 (magenta), and DAPI DNA stain (blue) with insets shown to the right (scale bar = 20 µm, 5 µm in inset), (representative example from n=3 biological replicates). A line-scan (yellow line) showing normalized fluorescent intensity of GFP (green) and GM130 (magenta) signal are plotted along the line (right). ***D***, Alignment of C-termini of human, *D. melanogaster*, and *C. elegans* DNAJC13/RME-8 with YLT1 motif highlighted in yellow (or the semiconserved region of *C. elegans* in orange).

**Figure S3.**
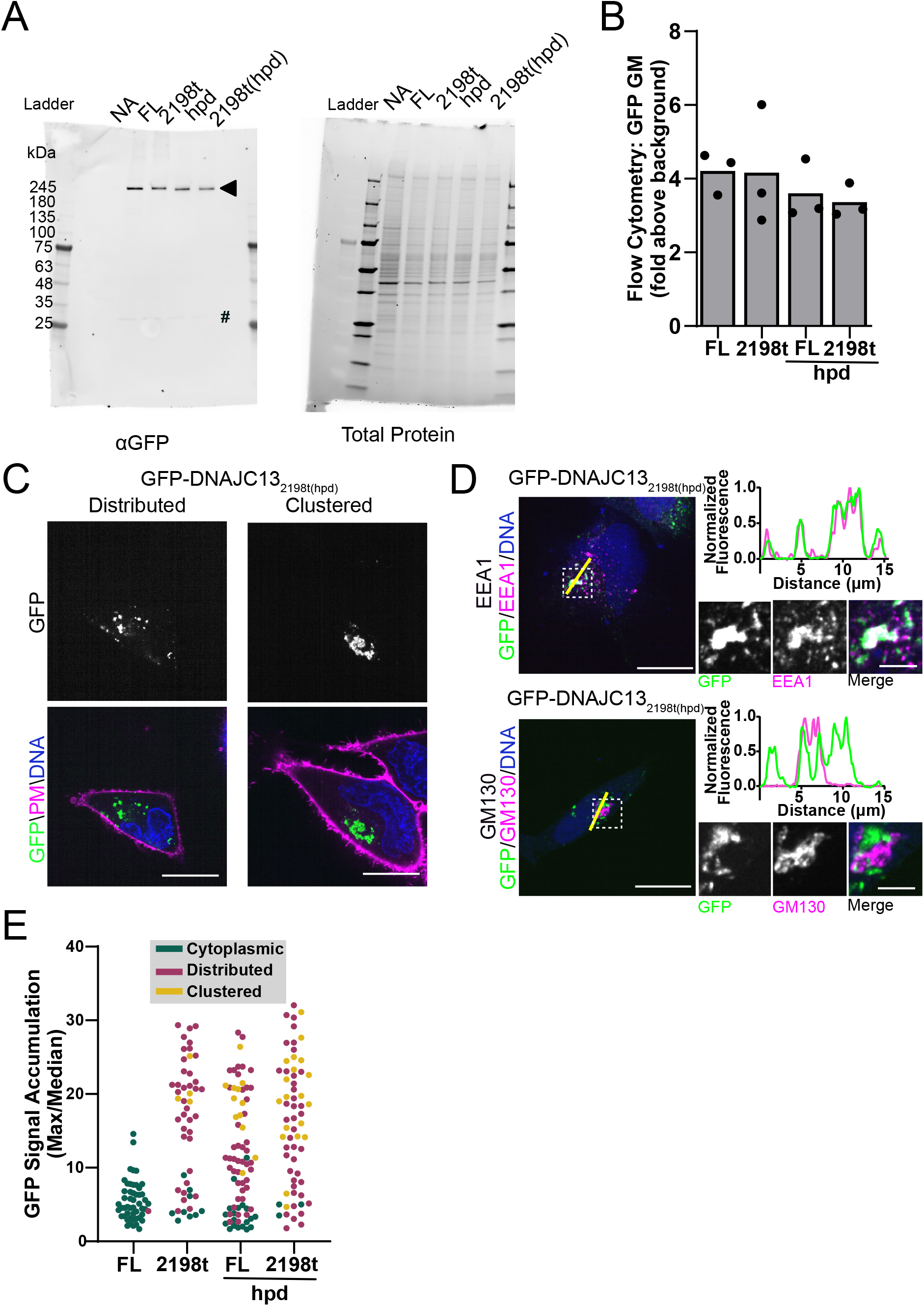
***A***, Representative western blot (anti-GFP, left) and total protein stain gel (right) of HeLa cells transfected with DNAJC13_FL_, DNAJC13_2198t_, DNAJC13hpd, or DNAJC13_2198t(hpd)_ and a nontransfected control in HeLa cells (n=3 biological replicates). The arrowhead marks GFP- DNAJC13, and the # marks free GFP. ***B***, Flow cytometry-based expression analysis of hpd mutant constructs in HeLa cells, assessed by geometric mean of GFP channel, displayed as fold above background signal from untransfected cells (n=3 biological replicates) . ***C***, Live cell spinning disk confocal microscopy images of GFP-DNAJC13_2198t(hpd)_ in HeLa cells showing distributed (left) and clustered (right) endosomes. Imaged with CellMask plasma membrane stain (magenta) and Hoechst DNA stain (blue) (scale bar = 20 µm) (phenotypic representative examples from n=3 biological replicates). ***D***, Fixed immunofluorescent microscopy image of GFP-DNAJC13_2198t(hpd)_ expressed in HeLa cells. Imaged with anti-GFP (Green), DAPI DNA stain (blue), and endosomal marker anti-EEA1 (magenta, top) or Golgi marker anti-GM130 (magenta, bottom). Insets shown to the right (scale bar = 20 µm, 5 µm in inset), (representative example from n=3 biological replicates). Line-scans (yellow lines) showing normalized fluorescent intensity of GFP (green) and EEA1 (magenta) or GM130 (magenta) signal are plotted along the lines (right). ***E***, Signal accumulation metric data (from Figure 3D), correlated by color to the blinded phenotypic scoring (from Figure 3E).

**Figure S4.**
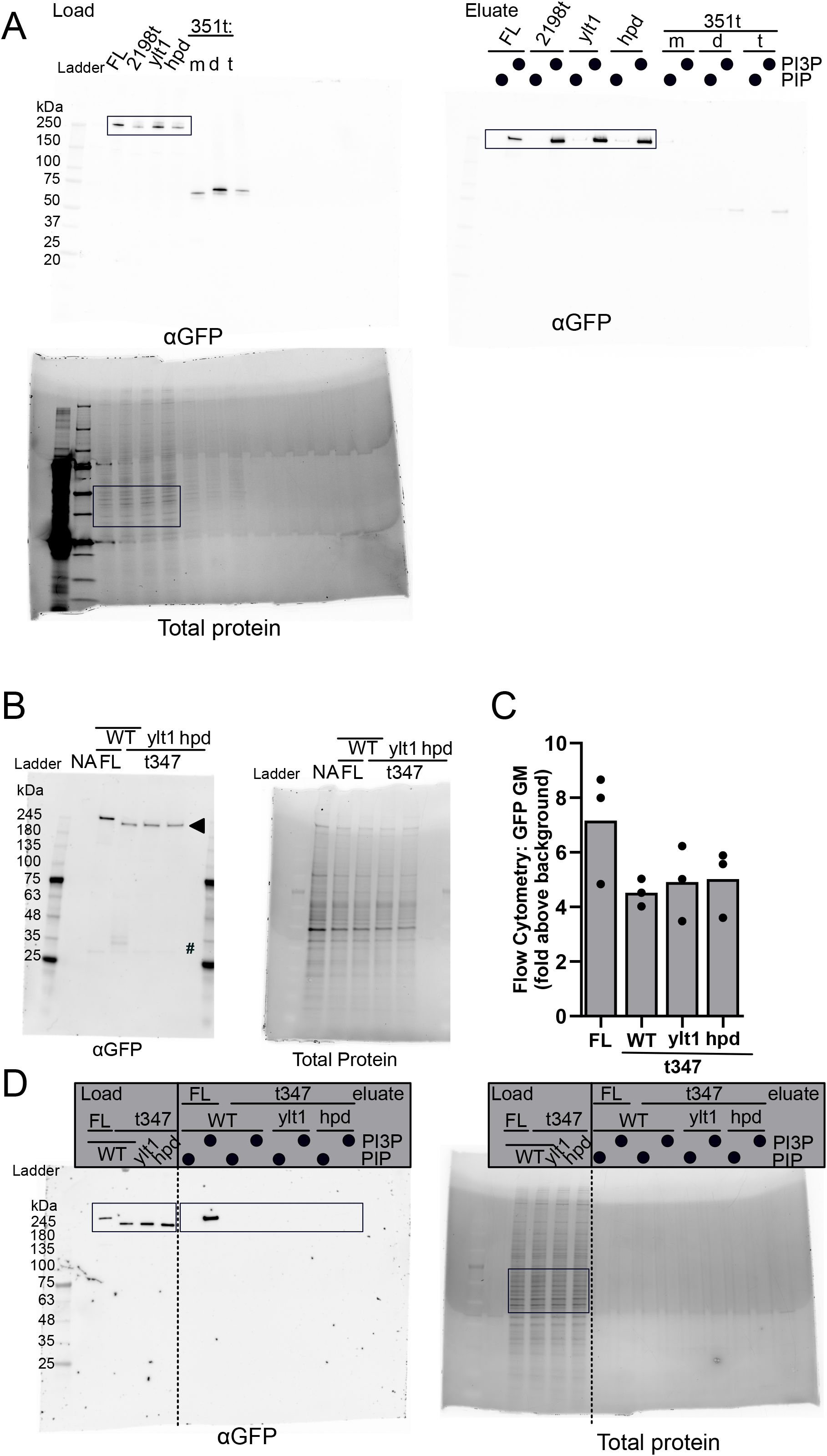
***A***, Uncropped blots (anti-GFP) and total protein stain gel from Figure 4A; cropped area shown in the black box. ***B***, Representative western blot (anti-GFP, left) and total protein stain gel (right) of HeLa cells transfected with DNAJC13_FL_, DNAJC13_t347_, DNAJC13_t347(ylt1)_, or DNAJC13_t347(hpd)_, and a nontransfected control (n=3 biological replicates). Arrowhead marks GFP-DNAJC13 and the # marks free GFP. ***C***, Flow cytometry-based expression analysis of t347 constructs in HeLa cells, assessed by geometric mean of GFP channel, displayed as fold above background signal from untransfected cells (n=3 biological replicates). ***D***, Uncropped blot (anti-GFP) and total protein stain gel from Figure 4C, cropped area shown in the black box.

**Figure S5.**
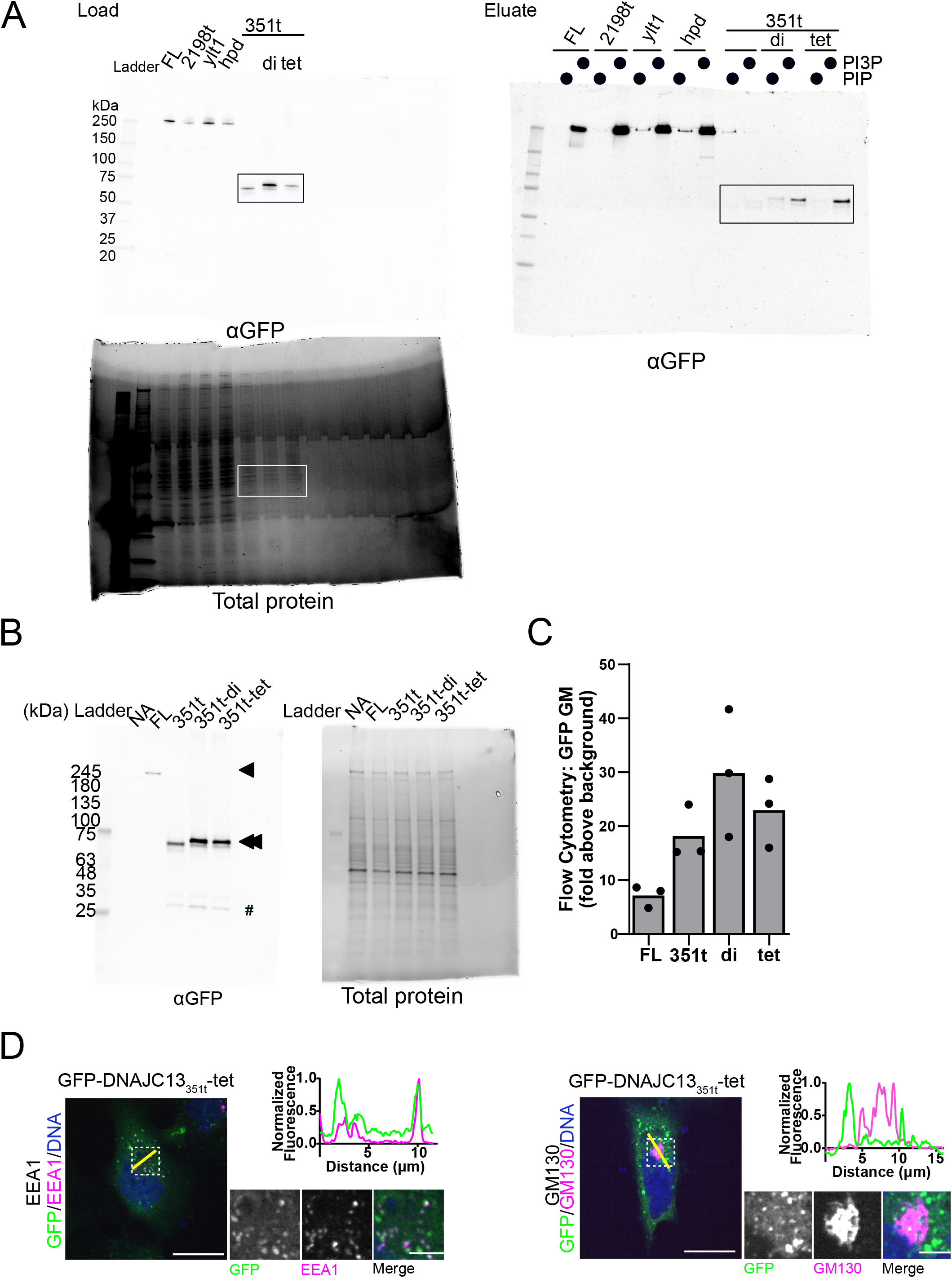
***A***, Uncropped blots (anti-GFP) and total protein stain gel from Figure 5A – gel was run with samples from Figure 4A, image was re-thresholded for viewing relevant samples, with cropped area shown in the black (or white) box. ***B***, Representative western blot (anti-GFP, left) and total protein stain gel (right) of HeLa cells transfected with DNAJC13_351t_, DNAJC13_351t_- dimer, or DNAJC13_351t_-tetramer, and a nontransfected control (n=3 biological replicates). Arrowhead marks GFP-DNAJC13_FL_, double arrowhead marks GPF-DNAJC13_351t_ and the # marks free GFP. ***C***, Flow cytometry-based expression analysis of DNAJC13_351t_ constructs in HeLa cells, assessed by geometric mean of GFP channel, displayed as fold above background signal from untransfected cells (n=3 biological replicates). ***D***, Fixed immunofluorescent microscopy image of GFP-DNAJC13_351t_-tetramer expressed in HeLa cells. Imaged with anti-GFP (Green), DAPI DNA stain (blue), and endosomal marker anti-EEA1 (magenta, left) or Golgi marker GM130 (magenta, right) with insets shown to the right (scale bar = 20 µm, 5 µm in inset), (representative example from n=3 biological replicates). Line-scans (yellow lines) showing normalized fluorescent intensity of GFP (green) and EEA1 (magenta) or GM130 (magenta) signal are plotted along the line (right).

